# Vegetation density is the main driver of insect species richness and diversity in small private urban front gardens

**DOI:** 10.1101/2024.06.12.598424

**Authors:** Joeri Morpurgo, Margot Huurdeman, J. Gerard B. Oostermeijer, Roy P. Remme

**Affiliations:** Department of Environmental Biology, Institute of Environmental Sciences, Leiden University, Einsteinweg 2, 2333 CC Leiden, Netherlands; Institute for Biodiversity and Ecosystem Dynamics, Department of Evolution and Population Biology, University of Amsterdam, Netherlands; Ministerie van Landbouw, Natuur en Voedselkwaliteit, The Hague, Netherlands

**Keywords:** Biodiversity, herbivores, nature-based solutions, pollinators, urban ecology

## Abstract

Urbanisation changes the natural ecosystems and vegetation to urban green spaces, and causes insect communities to experience novel challenges for survival. New evidence suggests that urban green spaces, no matter how small, can provide meaningful habitats for insects. Information on design and management of small gardens (<10m^2^) in dense urban areas is still scarce. In particular, it is hardly known which garden designs provide most benefits to insects. We surveyed 65 small private urban front gardens (μ=1.7m^2^) in Amsterdam and The Hague in The Netherlands and measured a series of garden attributes thought to be relevant for general, flower-visiting and herbivorous insect species richness and diversity. Plant coverage and richness were the strongest predictors for insect biodiversity and species richness. We found no support for associations with native vs. exotic plants or garden size.

**Synthesis and applications:** To strengthen insect biodiversity in the urban environment, we recommend future design of urban green spaces to focus on maximising coverage and richness of vegetation, potentially even using exotic species to fill in the gaps where native plant species cannot survive.

**Highlights:**

- Small urban gardens hold large potential for supporting urban insect communities
- Total vegetation cover was the strongest predictor for insect diversity and richness
- Plant richness was the second strongest predictor, but not for herbivorous insects
- Garden size had no effect on insect diversity or richness
- Native vegetation did not impact insect diversity or richness

Graphical abstract. Shown are either plus or minus signs indicating a positive or negative trends from the models that predict species richness and Shannon biodiversity of insects, pollinators or herbivores by the design of small urban gardens. ^*^indicates a statistically significant trend at p-value < 0.05. A plus or minus sign without^*^ means a visible statistically non-significant trend. A greyed out square indicates exclusion of the term in the model. A white square indicates non-significant and non-visible trend.

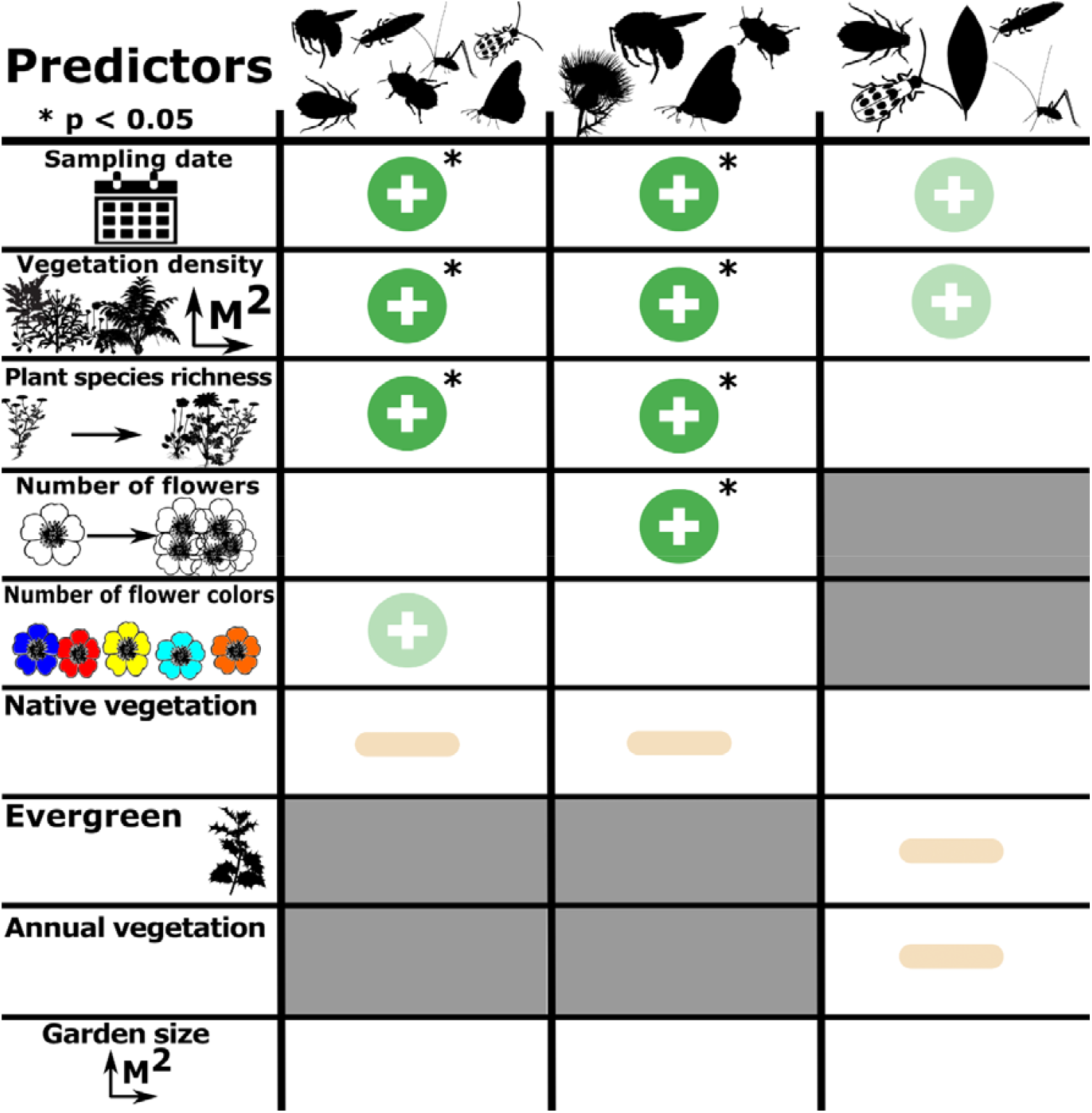

## 1. Introduction

Insects are essential for urban ecosystem functioning, health, and for supporting ecosystem services beneficial to humans. The economic value generated by insects is therefore often estimated to be worth billions (Losey & Vaughan, 2006; Ngo et al., 2017). In the urban environment, increasing insect biodiversity may support functions important to both humans and the ecosystem, such as pollination of flowers, nutrient cycling, supporting water regulation through soil-bioturbation, and as very important components of many trophic networks (Buchholz & Egerer, 2020; Belovsky and Slade, 2000; Drager et al., 2016; Eggleton, 2020). Flower-visiting insects are, for example, indispensable for pollination and therefore help sustain plant and insect populations throughout the urban and peri-urban environment (Eggleton, 2020). Herbivorous insects, sometimes considered pests by homeowners, are important to ecosystem functioning by making plant resources available for detritivores which in turn feed the soil. This trophic cascade from herbivorous insects to soil nutrients is highly important for plant diversity and ecosystem health (Belovsky and Slade, 2000). Despite the importance of insects to humans and ecosystems, their numbers are dwindling at alarming rates (Hallmann et al. 2017). With the demise of insects, many of the important ecosystem services we depend on, as well as biodiversity at large, will further destabilise to a point of no return (Wagner et al., 2021). Therefore, it is urgent to understand the drivers of insect diversity and provide meaningful information to policymakers and citizens so that we can support insects from an urban to global scale.

During urbanisation, changes to the natural landscape play a key part in reducing biodiversity (Güneralp & Seto, 2013). Here, urbanisation mostly affects natural vegetation through processes of fragmentation, degredation or complete removal. The novel urban green spaces (UGS) are more heterogeneous and differin composition from natural vegetation (Yang et al., 2021). As a consequence, the process of urbanisation has been shown to massively decrease in the abundance and species richness of several arthropod taxa, with butterflies showing the largest decline of 85% decrease in abundance (Piano et al., 2020)While initial urbanisation invokes the loss of local biodiversity, the new urban green spaces built are able to be colonized by species that are able to adapt to these different habitats and environmental conditions.

A key driver for biodiversity across the globe, and in cities, is the area of available habitat in urban areas (Beninde et al., 2016). Interestingly, the effect of UGS size on biodiversity has largely been ignored in urban ecological studies (Morpurgo et al., 2023). More importantly, at the relatively small spatial scale of privately owned gardens in urbanised areas (<10m^2^), size is to our knowledge rarely included or studied as a potentially important driver of biodiversity. Considering that a substantial amount of the UGS in cities are privately owned and small, they may contribute strongly to the urban biodiversity (Baker et al., 2018; Davies et al., 2009). For these reasons, we have focussed on private small urban front gardens (μ=1.7m^2^), which are small green strips directly in front of houses, often created by the inhabitants.

Some efforts have been made to understand the relationships between garden size and biodiversity in the urban setting (Majewska & Altizer, 2018). A recent extensive review of the literature showed that several studies especially link garden size to positive effects for plants biodiversity (Delahay et al., 2023). Garden size being important to insect diversity has also been confirmed by a meta-anlysis, yet importantly only includes studies on larger gardens (Majewska and Altizer, 2020; 6.1m^2^ – 70 000m^2^). The smallest garden sizes studies on insect species, were on urban grasslands using plots of flowering plants ranging from 1 to 100m^2^ and showing a positive correlation between area and insect pollinator diversity (Blaauw & Isaacs, 2014). In parallel, with increased urbanisation, the area of vegetation, and thus habitat, maybe a more mechanistic driver than garden size but this has not been properly investigated. In particular, the effect of plant coverage and vegetation layers on insect biodiversity have hitherto not been studied in small urban gardens.

In general, garden design is thought to be an important driver of urban insect biodiversity. A argued design choice is the inclusion ofnative plant species to provide greater benefits to biodiversity than exotic species (Schleapfler et al., 2011). The key mechanism explaining this benefit stems from a perceived mismatch between the traits of native fauna and exotic or ornamental plant species. This can for example lead to bees being unable to retrieve nectar from flowers that are shaped differently than native counterparts (Cohen et al., 2021). For herbivorous insects, exotic plant species may simply be inedible due to physical or biochemical resistance (Costan et al., 2020; Elton, 2020; i.e. enemy-release hypothesis). A study on floral resources and bee communities reported that, indeed, increasing proportions of native plants in gardens were associated with an increase in species richness and abundance of specialist bees (Tavares Brancher et al., 2023). Similarly, a study on backyard gardens reports that native plants in combination with local and landscape characteristics may be important for protecting and harbouring bee communities (Pardee & Philpott, 2014). However, there is a lack of evidence to support this narrative across the wider array of insect taxa. Salisbury et al. (2015) found that in addition to native species, some exotic species can play an important role in garden design by providing abundant resources to specialistic taxa and by extending the flowering season.

In this study, we aim to understand how the design of very small urban gardens (<10m^2^) is related to general, flower-visiting and herbivorous insect biodiversity. The ultimate goal is to identify easily adjustable design aspects of small gardens that increase insect diversity. A better understanding of how individual aspects of garden design impact insect diversity will better inform policymakers and citizens on how to design insect-friendly gardens. These may then more effectively support urban insect communities. To assess their impact, we measured several design attributes from private small gardens and related these to the insects we captured and identified. We modelled species richness and Shannon diversity separately for general, flower-visiting and herbivorous insects, using the garden attributes as predictors.

## 2. Methods

### 2.1 Selection of gardens

In 2019, we surveyed insects and vegetation of gardens (n = 65; Appendix 1) in the Dutch cities Amsterdam (52º21’33.1”N 4º54’33.5”E) and The Hague (52º04’57.0”N 4º17’40.3”E). These cities were chosen to exclude local confounding factors while they were also similar regarding the most relevant biotic and abiotic drivers. Climate is roughly the same, as the cities are 51km apart from each other, with a national average 851mm of precipitation and 10.5°C over the past 30 years (KNMI, 2023). Amsterdam is a larger city (21930ha) and The Hague is approximately half the size (9813ha). Both cities are classified as extremely urbanised, but The Hague has a higher population density of 6827 individuals per square kilometre than Amsterdam, with 4880 individuals per square kilometre (Centraal Bureau voor Statistiek, 2023).

To analyse the relationships between the attributes of small gardens and insects, we studied small private front gardens, as they are both abundant in both cities and small in size/area (<10m^2^). These gardens, façade garden, are defined as small gardens directly in front of a house, facing the public street. Prior to our field work, we asked the owners for permission to capture and examine invertebrates in their garden by distributing leaflets door-to-door. A façade garden was included in the study when (i) the homeowners granted permission, (ii) at least one plant was present in the garden, and (iii) the garden was situated within the spatial boundaries of the research area (Fig. 1, n = 65). This resulted in 50 sampling sites in Amsterdam and 15 in The Hague, which also generally had fewer façade gardens. To avoid spatial or temporal autocorrelation, we visited the sites in random order within and among cities.

**Figure 1.**
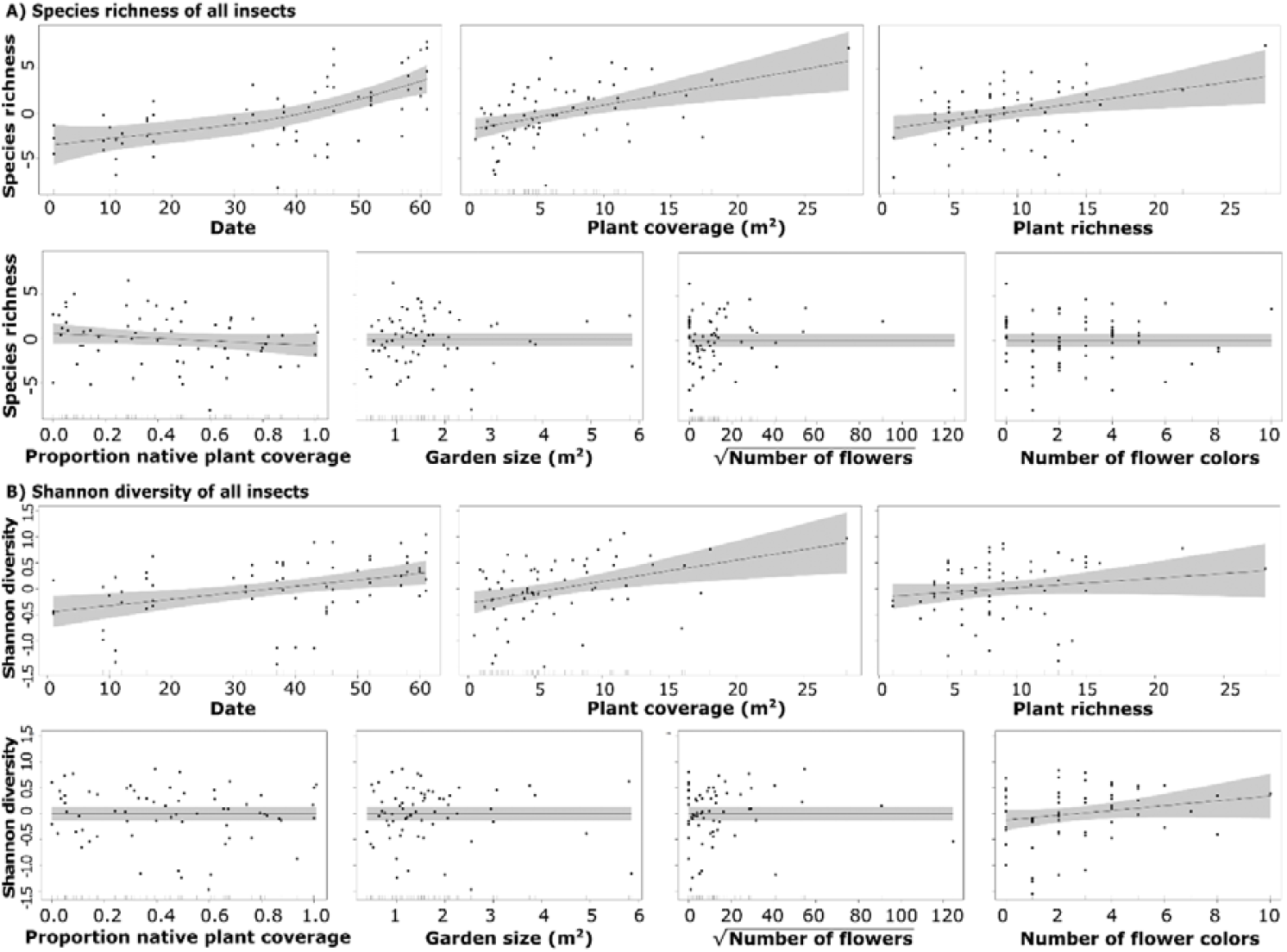
Smooths of seven predictor variables in the Generalized Additive Models predicting A) Species richness and B) Shannon diversity of all captured insects.

**Figure 2.**
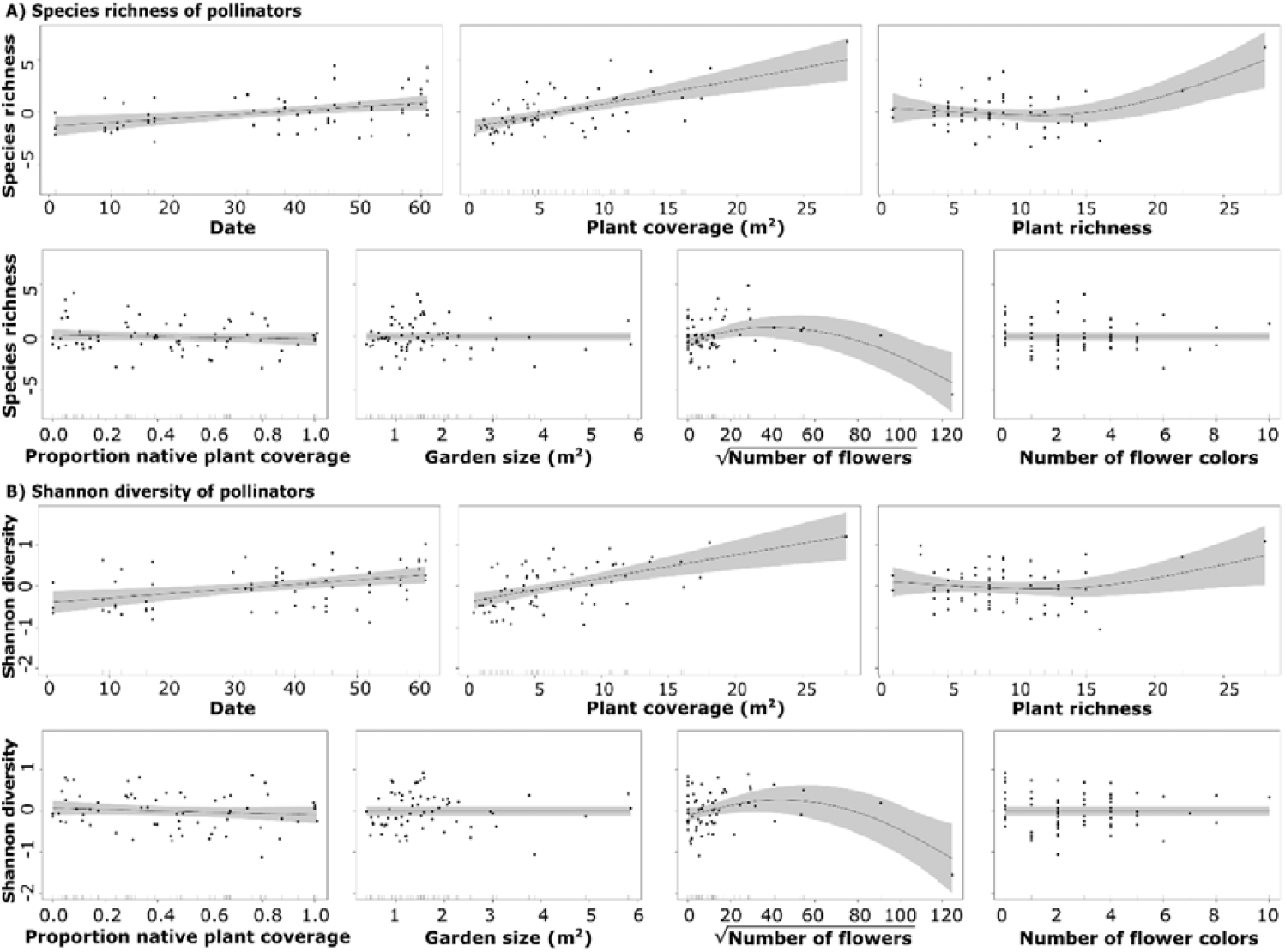
Smooths of seven predictor variables in the Generalized Additive Models predicting A) Species richness and B) Shannon diversity of flower-visiting insects.

**Figure 3.**
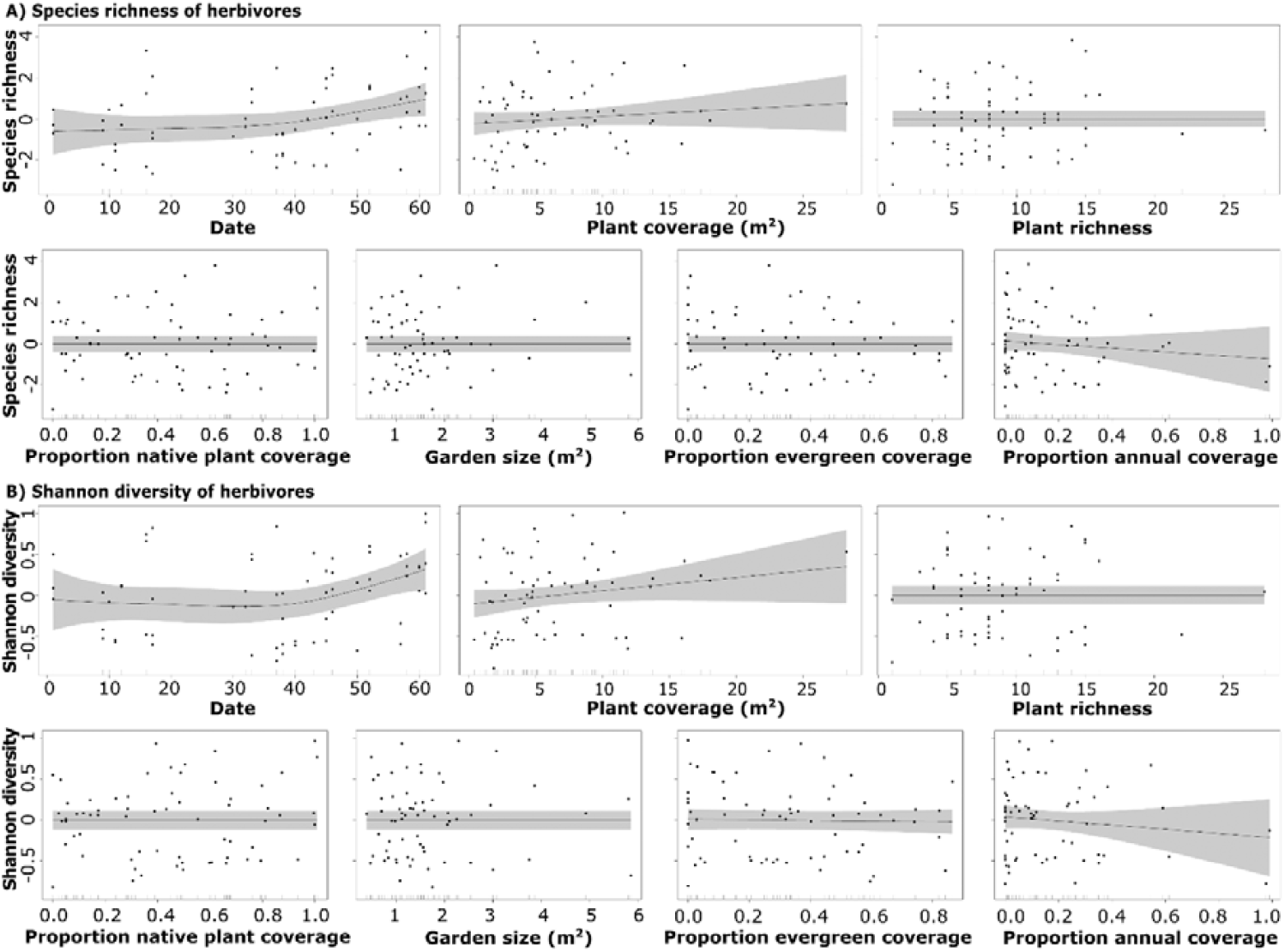
Smooths of seven predictor variables in the Generalized Additive Models predicting A) Species richness and B) Shannon diversity of herbivorous insects only.

### 2.2 Design of the gardens

We collected data on several features of each garden and its environment that might affect insect diversity (Table 1). The cardinal direction of the garden was included, as it impacts the amount of sunlight received. Subsequently, we recorded the temperature, time of survey, date, and the area in square metres. Within the garden, temperature was read from a thermometer in a spot illuminated by the sun, with exception of gardens that were in the shade during sampling.

**Table 1.**
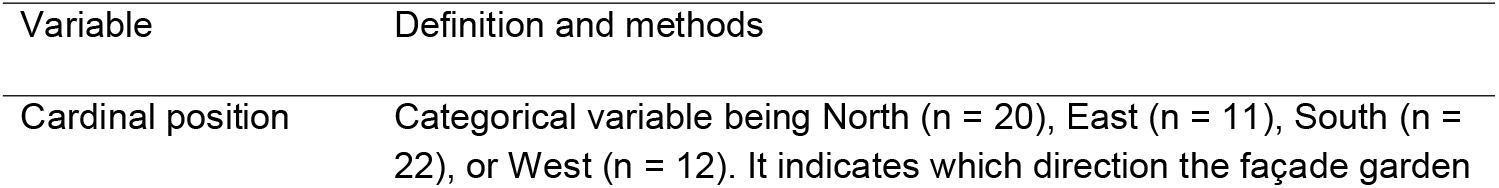
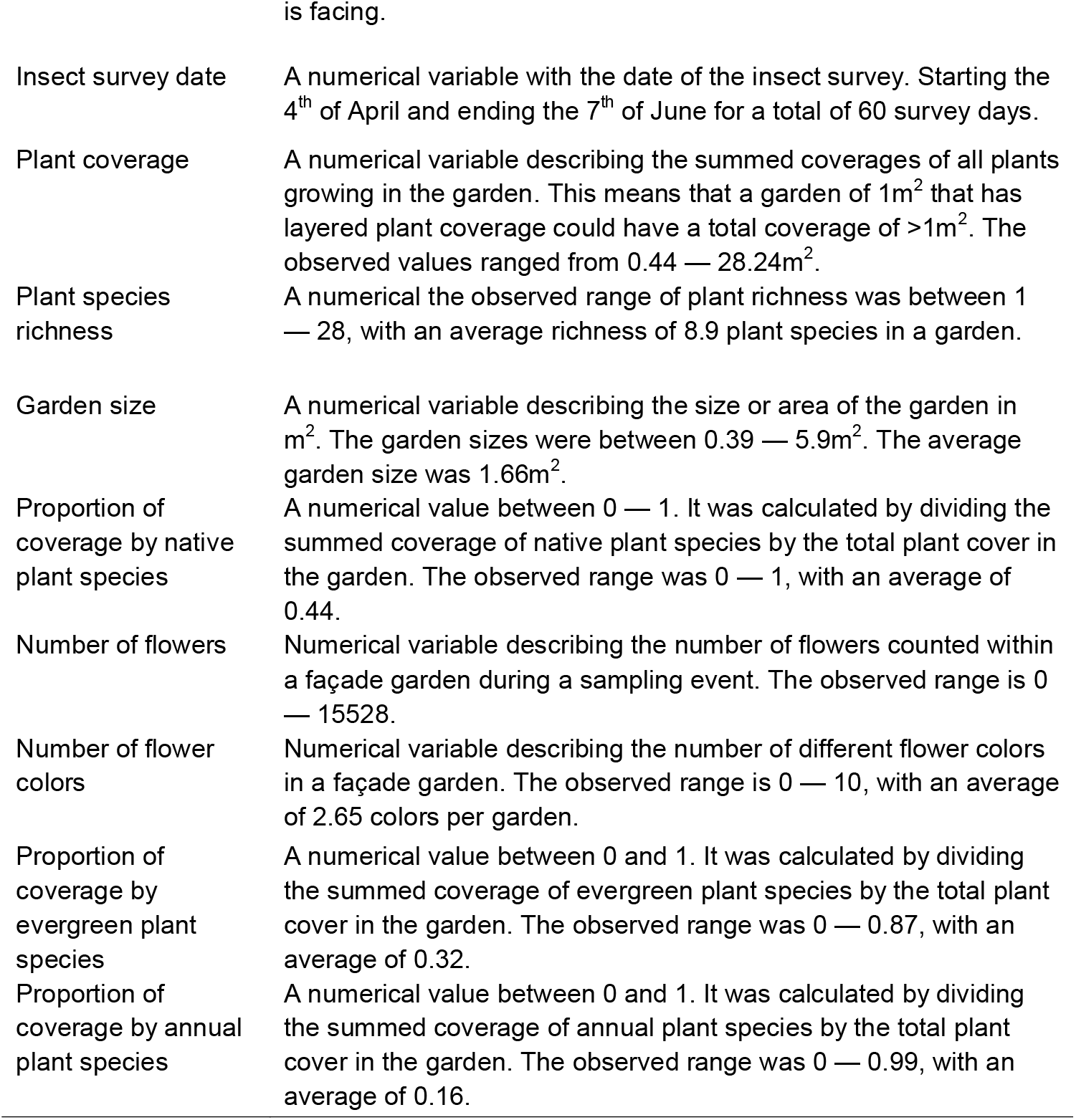
Summary of variables sampled thought to explain insect abundance and diversity (response variables). The predictor variables that capture attributes of small gardens (size, flowers, litter), climate, and temporality.

### 2.3 Vegetation surveys

Vegetation surveys were conducted before and after the insect survey. All plant species were identified and classified as native using ‘Heukels’ Flora van Nederland (Heukels and Van der Meijden, 2005). Species absent from the ‘Heukels’, such as several cultivars or exotic species, were identified using Pl@ntNet (Affouard et al., 2023).

For each individual plant species, we measured both cover and number of flowers (Table 1). For every species, we measured total coverage by holding a tapeline crosswise aiming to maximising the coverage of the individual and summing the coverages of all individuals. The number of flowers per plant was counted if it had fewer than 100 flowers, and for plants with more than 100 flowers we counted the number on a single representative flower cluster and multiplied it by the total number of clusters. Finally, we also counted the number of easily distinguishable flower colors visible to the human eye (e.g. red, blue, but also dark blue or light blue).

### 2.4 Insect survey

Insect surveys were performed from April until June 2019 between 09:00 and 17:00, excluding only rainy days. Surveys were based on sampling efforts of 15 minutes, during which insects were thoroughly searched on the aboveground parts of plants by two researchers simultaneously. Observed insects were captured using 50 ml falcon tubes and visually identified after the survey. Species were directly identified in the field using Chinery (2007) and released after the survey. Unidentified species were euthanized with ethyl acetate and identified in the lab using additional literature and ObsIdentify (Albouy, Richard, 2019; Ball and Morris, 2015; Van Veen, 2010). Finally, if a species could not be identified, it was assigned to a morphotype comprising morphologically identical organisms.

### 2.5 Statistical analysis

We employed six separate statistical models to link garden design features to aboveground insect diversity. To investigate the relationships more comprehensively, we created separate models predicting either species richness or Shannon diversity for: I) all observed insects, II) only flower-visiting insects, and III) only herbivorous insects. These categories were made to infer if all observed insects were affected in the same way, or whether flower-visiting or herbivorous taxa are affected by different garden features. We chose to use Generalised Additive Models (GAM) as they allow for non-linear relationships accounting for seasonality of insect richness and diversity, while favouring linear ones (Wood et al., 2017).

Considering the above, the mathematical formulation of the GAM for all insects and only flower-visiting insects was:

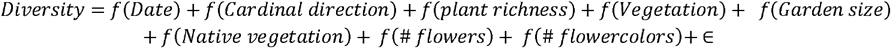

For herbivores, we changed the terms that are flower-related to leaf-related traits. Therefore, the GAM for insect herbivores is:

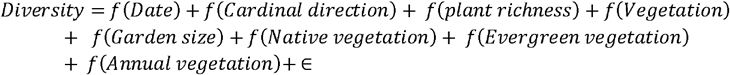

In all models, *diversity* represents either species richness or Shannon diversity index of either all insects captured, or solely flower-visiting or herbivorous insects. The term *Date* used a Gaussian Process smoothing that accounts for temporal autocorrelation, and relates to the number of days since the first insect survey. The *Cardinal direction* term uses a random effect to account for average differences in insect richness or diversity for gardens which face different directions, with North-facing gardens receiving least, and South-facing ones most sunlight. Finally, ∈ represents the unexplained error term of the model.

The other predictors are all modelled smooths using thin plate regression splines (TPRS) allowing for non-linear relationships (Wood, 2003). These splines effectively act as a model fitting step as the null space is also penalized slightly and the whole term can therefore be shrunk to zero (Wood, 2003). During the model fitting, several knots are used to smooth linear regression to a spline optimizing the restricted maximum likelihood (REML; Wood, 2017, p. 185). Effectively, this means that zero knots equals a linear regression, and every additional knot added allows for more smoothing. The advantage of the used TPRS approach is that knot number and position are based on the data instead of manually and subjectively placing knots.

The terms that used the TPRS smooths are I) *vegetation* as it relates to the summed coverage of every species in a garden, II) *native vegetation* which is a proportion of the summed native plant coverage in a garden, compared to that of exotic species, III) the *number of flowers*, and IV) *the number of flower colors* terms, which relate to the number of flowers and the number of visible flower colors in a garden, V) *evergreen vegetation*, and VI) *annual vegetation* which both relate to the proportion of the summed coverage of the respective vegetation component.

The model diagnostics were visually inspected through assessing the autocorrelation function of the residuals, concurvity and the model residuals. No issues of concurvity (i.e. the non-linear variant of multicollinearity) or model fit were identified (Appendix 3 — 8).

## 3. Results

### 3.1 Data collection

Across the 65 observed façade gardens (Amsterdam (n = 50), The Hague (n = 15)) we found 235 plant and 154 insect species. The gardens had an average area of 1.7m^2^, while the largest garden comprised 5.9m^2^. Of all plant species identified, 40% were native (n = 94), 26% were evergreen (n = 61), and 17% were annual (n = 39). Plant species richness ranged from 1 – 28 with a mean of 8.9 species per garden. Of all the insect species, 55% were identified as a flower-visitor (and potential pollinator, n = 85), and 25% as a herbivore (n = 39). The insect species richness observed in the gardens ranged from 0 — 19 with a mean of 8 species per garden.

### 3.2 GAM for all insects

The GAMs explained the data containing all the insects with a deviance of 57.8% for species richness (R^2^ = 0.54; Fig. 1A) and 40.0% for Shannon diversity (R^2^ = 0.37; Fig. 1B). In both models, sampling date and plant coverage were significantly positive predictors (p < 0.01; Table 2). Plant richness was positively significant in the insect species richness model (p < 0.01). The number of flowers, the number of flower colours, the proportion coverage by native plants and garden size were not significant in either model (p > 0.10).

**Table 2.** Results of Generalized Additive Models (GAMs) to explain species richness and Shannon diversity of all insects captured in façade gardens (n = 65). The intercept is shown with parametric coefficients, whereas covariates are represented with smooth terms. For details of predictor variables see table 1. Significant *p*-values (*p* < 0.05) are indicated in bold. *edf – Estimated Degrees of Freedom

| All insects | Species richness GAM |  |  |  | Shannon diversity GAM |  |  |  |
| --- | --- | --- | --- | --- | --- | --- | --- | --- |
| Predictors | *Estimate* | *Std. error* | *t-value* | *p-value* | *Estimate* | *Std. error* | *t-value* | *p-value* |
| *Intercept* | 8.01 | 0.39 | 20.55 | **<0.001** | 1.5 | 0.07 | 22.6 | **<0.001** |
| Smooth terms |  | *edf\** | *F-value* | *p-value* |  | *edf\** | *F-value* | *p-value* |
| *f (Cardinal direction)* |  | 0.41 | 0.16 | 0.31 |  | <0.001 | 0 | 0.87 |
| *f (Date)* |  | 2.06 | 17.15 | **<0.001** |  | 1.00 | 10.83 | **0.002** |
| *f (Plant richness)* |  | 0.91 | 0.89 | **0.006** |  | 0.67 | 0.22 | 0.08 |
| *f (Vegetation coverage)* |  | 0.98 | 1.59 | **<0.001** |  | 0.95 | 1.15 | **0.001** |
| *f (Proportion of native vegetation)* |  | 0.65 | 0.21 | 0.10 |  | <0.001 | 0 | 0.41 |
| *f (Number of flowers)* |  | <0.001 | 0 | 0.78 |  | <0.001 | 0 | 0.67 |
| *f (Number of flower colors)* |  | <0.001 | 0 | 0.70 |  | 0.73 | 0.31 | 0.051 |
| *f (Garden size)* |  | <0.001 | 0 | 0.99 |  | <0.001 | 0 | 0.67 |
| Adjusted R<sup>2</sup> | 0.54 |  |  |  | 0.37 |  |  |  |
| Explained deviance | 57.8% |  |  |  | 40% |  |  |  |

### 3.3 GAM on flower visitors

The GAMs explained the data of only flower-visiting insects very well, with a deviance of 63.3% for species richness (R^2^ = 0.59) and 56.8% for Shannon diversity (R^2^ = 0.52). In both models, sampling date and plant coverage contributed significantly positive (p < 0.01; Table 3). For both species richness and Shannon diversity, the number of flowers showed a significant curvilinear relationship (p < 0.01). Flower visiting species richness also correlated with plant species richness (p < 0.01). In contrast, neither model showed a significant contribution of the number of flower colors (p = 0.64), and the same was true for the proportion of coverage by native plants and garden size (p > 0.10).

**Table 3.** Results of Generalized Additive Models (GAMs) to explain species richness and Shannon diversity of flower-visiting insects in façade gardens (n = 65). The intercept is shown with parametric coefficients, whereas covariates are represented with smooth terms. For details of predictor variables see table 1. Significant *p*-values (*p* < 0.05) are indicated in bold. *edf – Estimated Degrees of Freedom

| Flower-visiting insects | Species richness GAM |  |  |  | Shannon diversity GAM |  |  |  |
| --- | --- | --- | --- | --- | --- | --- | --- | --- |
| Predictors | *Estimate* | *Std. error* | *t-value* | *p-value* | *Estimate* | *Std. error* | *t-value* | *p-value* |
| *Intercept* | 2.98 | 0.19 | 15.32 | **<0.001** | 0.85 | 0.06 | 14.9 | **<0.001** |
| Smooth terms |  | *edf\** | *F-value* | *p-value* |  | *edf* | *F-value* | *p-value* |
| *f* (*Cardinal direction*) |  | <0.001 | 0 | 0.67 |  | <0.001 | 0 | 0.45 |
| *f* (*Date*) |  | 1 | 9.66 | **0.003** |  | 1.00 | 10.11 | **0.002** |
| *f* (*Plant richness*) |  | 2.51 | 1.59 | **0.001** |  | 1.94 | 0.59 | 0.06 |
| *f* (*Vegetation coverage*) |  | 1.07 | 3.02 | **<0.001** |  | 1.03 | 2.28 | **<0.001** |
| *f* (*Proportion of native vegetation*) |  | 0.40 | 0.07 | 0.19 |  | 0.55 | 0.14 | 0.14 |
| *f* (*Number of flowers*) |  | 2.36 | 1.60 | **0.001** |  | 2.30 | 1.35 | **0.003** |
| *f* (*Number of flower colors*) |  | <0.001 | 0 | 0.90 |  | <0.001 | 0 | 0.94 |
| *f* (*Garden size*) |  | <0.001 | 0 | 0.58 |  | <0.001 | 0 | 0.84 |
| Adjusted R<sup>2</sup> | 0.59 |  |  |  | 0.52 |  |  |  |
| Explained deviance | 63.3% |  |  |  | 56.8% |  |  |  |

### 3.4 GAM herbivores

The GAMs explained the data of only herbivorous insects comparatively poorly, with a deviance of 20.0% for species richness (R^2^=0.14) and 19.5% for Shannon diversity (R^2^=0.14). This is reflected in the lack of any smooth being significantly linked to richness or diversity of herbivores. The model did not show any significant relationship with a vegetation indicator (p > 0.05; Table 4). In both models, plant coverage showed visually the strongest relationship and was nearly significant for Shannon diversity (p = 0.07) and less for the species richness (p = 0.13).

**Table 4.** Results of Generalized Additive Models (GAMs) to explain species richness and Shannon diversity of herbivorous insects in façade gardens (n = 65). The intercept is shown with parametric coefficients, whereas covariates are represented with smooth terms. For details of predictor variables see table 1. Significant p-values (p < 0.05) are indicated in bold. *edf – Estimated Degrees of Freedom.

| Herbivorous insects | Species richness GAM |  |  |  | Shannon diversity GAM |  |  |  |
| --- | --- | --- | --- | --- | --- | --- | --- | --- |
| Predictors | *Estimate* | *Std. error* | *t-value* | *p-value* | *Estimate* | *Std. error* | *t-value* | *p-value* |
| *Intercept* | 2.47 | 0.24 | 10.27 | **<0.001** | 0.63 | 0.06 | 10.96 | **<0.001** |
| Smooth terms |  | *edf\** | *F-value* | *p-value* |  | *edf* | *F-value* | *p-value* |
| *f (Cardinal direction)* |  | 0.95 | 0.52 | 0.19 |  | <0.001 | 0 | 0.49 |
| *f (Date)* |  | 1.86 | 3.63 | 0.11 |  | 2.30 | 3.11 | 0.08 |
| *f (Plant richness)* |  | <0.001 | 0 | 0.80 |  | <0.001 | 0 | 0.91 |
| *f (Vegetation coverage)* |  | 0.59 | 0.16 | 0.13 |  | 0.74 | 0.30 | 0.07 |
| *f (Proportion of native vegetation)* |  | <0.001 | 0 | 0.93 |  | <0.001 | 0 | 0.88 |
| *f (Proportion of evergreens)* |  | <0.001 | 0 | 0.59 |  | 0.13 | 0.02 | 0.72 |
| *f (Proportion of annuals)* |  | 1.00 | 0.92 | 0.34 |  | 1.02 | 0.86 | 0.37 |
| *f (Garden Size)* |  | <0.001 | 0 | 0.62 |  | <0.001 | 0 | 0.92 |
| Adjusted R<sup>2</sup> | 0.14 |  |  |  | 0.14 |  |  |  |
| Explained deviance | 20% |  |  |  | 19.5% |  |  |  |

## 4. Discussion

### 4.1 Interpretation of results

Our results show that several design attributes of small private front gardens (μ=1.7m^2^) in urban areas are correlated with insect biodiversity. We observed a large diversity of 235 plant and 154 insect species in an area of just 108m^2^ across a total of 65 gardens. Most importantly, we show that the strongest predictor for insect species richness and biodiversity is the total amount of plant coverage and richness of a garden. Moreover, the results suggest that flower-visiting insects are most strongly dependent on the amount of vegetation and to a lesser extent on the availability of flowers. Such correlations are much less evident for herbivorous insects. Our results on the relationship between plant cover and richness driving insect diversity are generally congruent with previous research and suggest that even in small privately owned urban gardens, adding more plants and more plant species is quite likely to have a positive impact on the insect community (Pardee & Philpott, 2014; Beninde et al., 2015; Blaauw & Isaacs, 2014).

Interestingly, we find that flower-visiting and herbivorous insects responded quite differently to the design of façade gardens. As expected, flower visitors responded favourably to flower-related attributes (flower number and colour diversity) as they provide food, which is in line with previous studies (Quistberg et al., 2016). Overall, we found fewer species of herbivores than flower visitors in the façade gardens. Possibly, this is because flower-visiting insects are easier to detect than herbivorous insects which are generally hidden to avoid predation. Especially during the 15 minutes of capturing insects, flower-foraging insects are much more likely to fly from one garden to another to forage. Both the camouflage of herbivores and flying behaviour of flower-visiting insects would explain the result in a higher capture rate of flower visitors than of herbivores.

In contrast to our expectations, flower-visiting insects showed a significantly positive relation with vegetation cover whereas herbivorous insects did not. We propose three different mechanisms, which each may play a part in the different relationship with vegetation cover between herbivorous and pollinating insects. The first conceivable mechanism involves food availability, as herbivorous and flower-visiting insects have distinctly different food sources. Herbivores forage on leaves and stems, which the majority of plant biomass. While, flower visitors forage on the much scarcer flowers, requiring them to forage on a larger diversity of plants in larger areas. The second mechanism could be simply that more vegetation coverage does not necessarily equal to more food availability for herbivorous insects. For example, it has been demonstrated that pollinators have a more generalist food network compared to herbivores, which may result in a narrow selection of plants to feed off (Fontaine et al., 2009). This would result in herbivorous insects being more attracted to specific plant species, than pollinators attraction to a general community of plants (Underwood et al., 2020). A third mechanism may be that pesticides impact herbivorous species much more severely than flower visitors. Urban gardens are often designed with the intent to be either aesthetic, native or biodiverse, through rigorous design and management of the constituting (ornamental) plants (Noe et al., 2021). Gardens with high plant coverage may indicate a stronger intent from the owners to facilitate other species as well. Unfortunately, many owners regard herbivorous insects as pests and use pesticides to protect their garden, potentially resulting in a lower diversity of herbivorous insects. Many ornamental plants are even treated with pesticides before they are sold (Lentola et al. 2017). There is evidence that most pesticides used would also harm pollinators through different pathways than intended (Cecala et al. 2021; Chagnon et al., 2015; Sánchez-Bayo, 2021). Hence, the difference between flower-visitors and herbivores in our results could be partially explained by both differences in foraging strategies and the potential use of pesticides (Jones and Agrawal, 2017; Lowe et al., 2019).

Surprisingly, there was no relationship at all between the proportion of coverage by native plants in the garden and insect diversity or richness. Generally, native plants are assumed to be better for biodiversity (Fleishman et al., 2011; Berthon et al., 2021). Mostly because insects and plants originating from the same region are thought to have co-evolved and are therefore assumed to have innate interactions leading to resources otherwise deemed inaccessible for exotic insect species. More recently there seems to be a paradigm shift away from nativeness and towards species’ function and traits to explain biodiversity patterns (Davis et al., 2011). For example, some evidence suggests that canopy coverage is the most important driver for soil-surface and plant-associated invertebrates diversity. And, the effect of nativeness of plant species seems to be dependent on the invertebrate taxa studied and the traits or functions of the plant species (Davis et al., 2011; Salisbury et al., 2020).

We propose four possible mechanisms to explain the lack of a significant relationship between insect diversity and plant nativeness. A first explanation could be that the exotic plants have similar traits or functions to their native counterparts. Therefore, barriers between insect plant interaction on the basis of exotic status, is mitigated by matching the traits of native plants and insects. Another explanation could be that insect species in the urban environment, which are often generalists, may simply do not exhibit a specialistic relationship with floral traits. Therefore, an increase of exotic plant species, with different traits, may not limit insect plant interactions. Alternatively, insect species that are specialists and may require native plants may simply not be able to reach the gardens within a highly urbanised area through other constraints posed on them by the urban environment instead of by the vegetation itself (Catzim et al., 2022). For example, native species may simply not reach urban gardens if they are not able to disperse through the hotter and drier climate of a city. A final explanation could be that the majority of the found insect species may be exotic of origin. In this case, exotic insects are not co-evolved with the native plants and therefore do not benefit from trait matching. This could imply that exotic insects may not be able to forage food from native plants as effectively, making them unattractive to neutral gardens to the potentially exotic urban insect community. One or a combination of these mechanisms may explain why there is no apparent relationship between insect diversity and the proportion of native plant coverage in urban façade gardens.

### 4.2 Implications for management and research

This is one of the first studies focussing specifically on very small urban façade gardens. Our results show that, although small, façade gardens are capable of supporting insect communities. The results also show that to support and promote biodiversity in urban green spaces, it may be more effective to focus on increasing overall plant coverage and richness rather than specifically promoting native plant species. City managers and urban gardeners could consider incorporating non-native plants strategically, such as using them to fill gaps in plant coverage where native species may not have the required traits to survive in the harsh urban environment. Many non-native plants are very attractive to (generalist) insect flower-visitors, and may increase their resources and consequently population sizes (Salisbury et al. 2015). These non-native plants could still provide ample benefits to biodiversity in contrast to the other dominant options that the urban environment provides, namely concrete or asphalt. While native plants may not be the most important predictor of insect biodiversity in small urban gardens, they can still play a valuable role in supporting ecosystems and provide meaningful services. For example, native plants have been shown to provide substantial cultural benefits in sacred sites (Gopal et al., 2017). Also, native plants may maintain the populations of their insect partner in the native species-specific relationship (Mitchell et al., 2009). Future scientific endeavours should focus on understanding the links between plant and insect communities along the urban gradient. One of the more pressing questions is whether the relationship found between summed vegetation coverage and insects is moderated by urbanisation. Which factor is more limiting to a thriving insect community: the lack of vegetation or increased urbanization? And at what level of urbanisation does habitat become a limiting factor to insects? Additionally, we must further acknowledge a lack of understanding of which type of species are impacted more by a shortage of suitable habitat. For example, specialist species may be more vulnerable to a species-poor plant community than generalist species. The challenge of finding the appropriate food could also be amplified by a species’ capacity to forage over longer distances.

## 5. Conclusions

Protecting biodiversity across urban areas is crucial to meeting targets set by global conservation frameworks (e.g., Sustainable Development Goals, Intergovernmental Science-Policy Platform on Biodiversity and Ecosystem Services and the Global Biodiversity Framework). The abundant private small urban green spaces and their design are important to enhancing insect biodiversity. The investigated gardens showed a tremendous variety in plant (n = 235) and insect species (n = 154). Our analysis shows that the design of very small gardens (μ=1.7m^2^) matters, and that increasing vegetation density and plant richness increases insect diversity. Based on our results, an extensive network of small gardens with high vegetation density may prove as valuable as a single large urban green space such as a city park. We recommend that policymakers and practitioners focus on promoting non-destructive gardening practices, aiming to increase the density and richness of vegetation with urban gardens, in order to support and strengthen insect communities. Future research should focus on delineating a potential moderating effect of urbanisation on different types of species.

## Supporting information

Map

Herbivore models

Pollinator models

Insect models

## 6. Data accessibility

Collected data, search queries and code used for this study have been made permanently and publicly available on the Mendeley Data repository at TBA.

## List of appendices

Appendix 1. Maps with boundaries of the sampling location

Appendix 3. Model diagnostic plots from the GAM predicting pollinator species richness

Appendix 4. Model diagnostic plots from the GAM predicting pollinator diversity

Appendix 5. Model diagnostic plots from the GAM predicting herbivore species richness

Appendix 6. Model diagnostic plots from the GAM predicting herbivore diversity

Appendix 7. Model diagnostic plots from the GAM predicting insect species richness

Appendix 8. Model diagnostic plots from the GAM predicting insect diversity

