## Supplementary material for "Vegetation density is the main driver of insect species richness and diversity in small private urban front gardens": Map

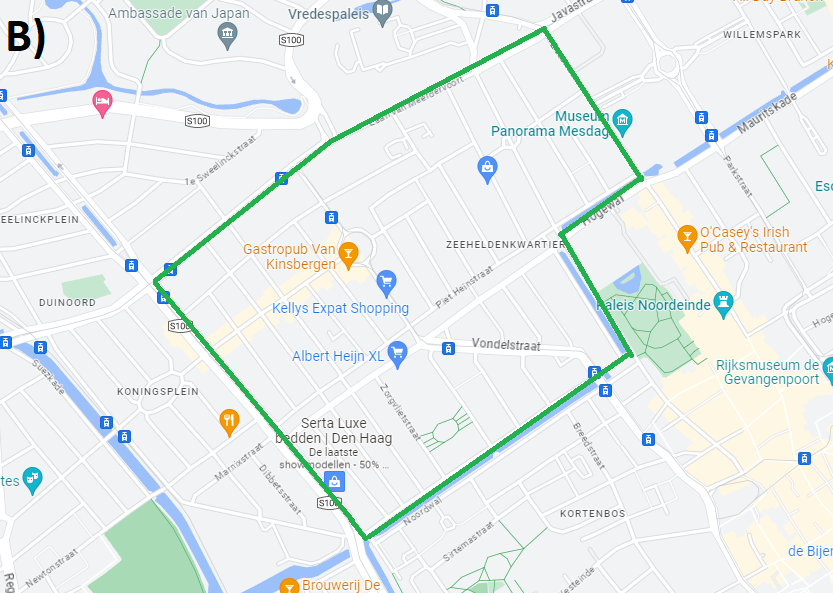

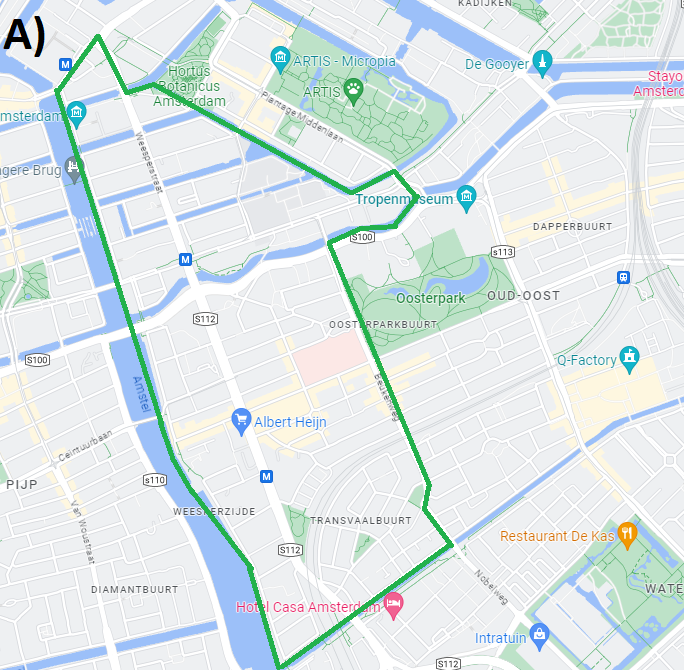

**Appendix 1. Shown are the boundaries of the sampling locations in green.** The locations within the cities of A) Amsterdam and B) The Hague are generally of similar size. Interestingly, the Amsterdam location has several parks close to the sampling location. The Hague has several larger parks slightly further from the sampling location.
