## Supplementary material for "Vegetation density is the main driver of insect species richness and diversity in small private urban front gardens": Herbivore models

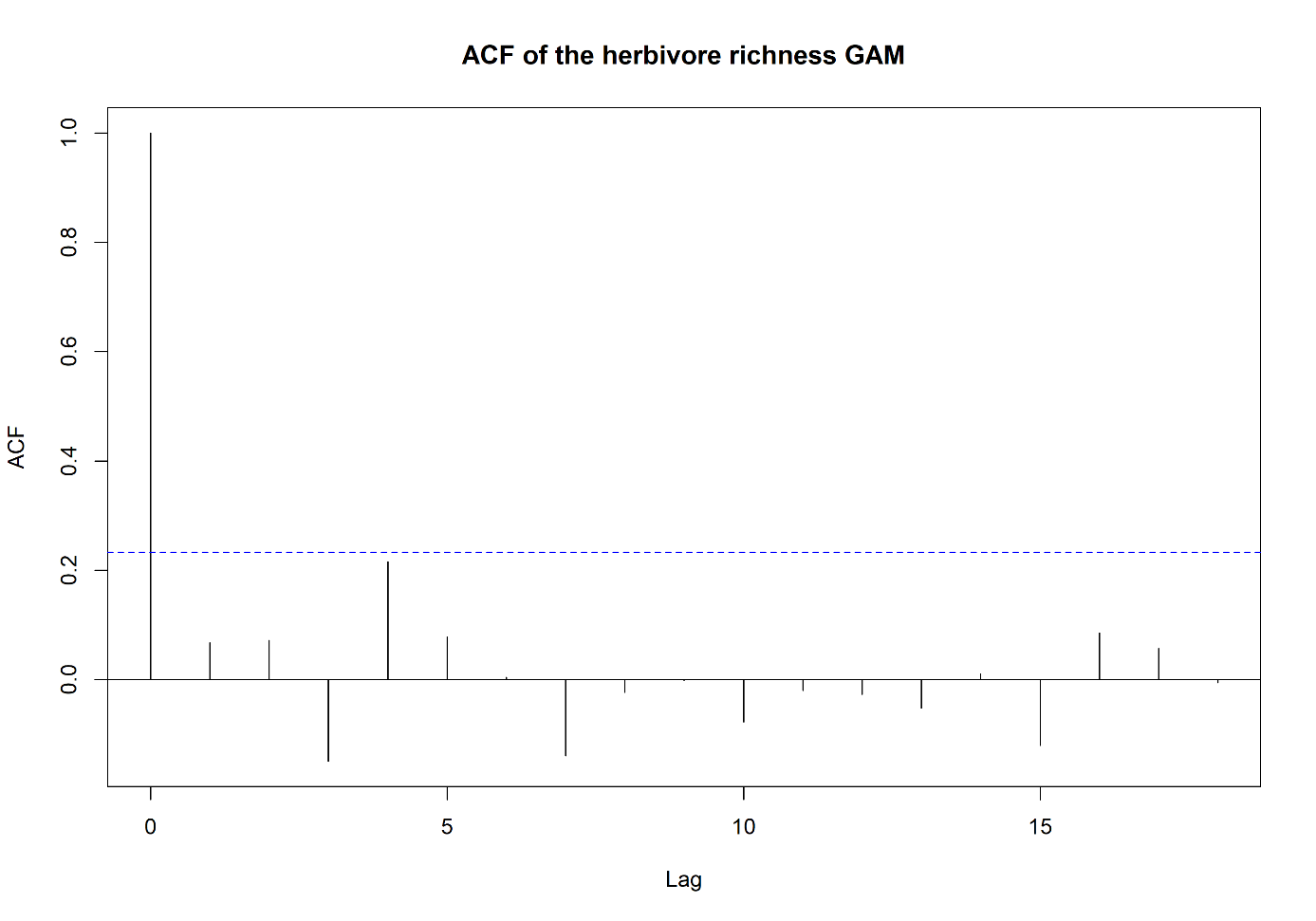

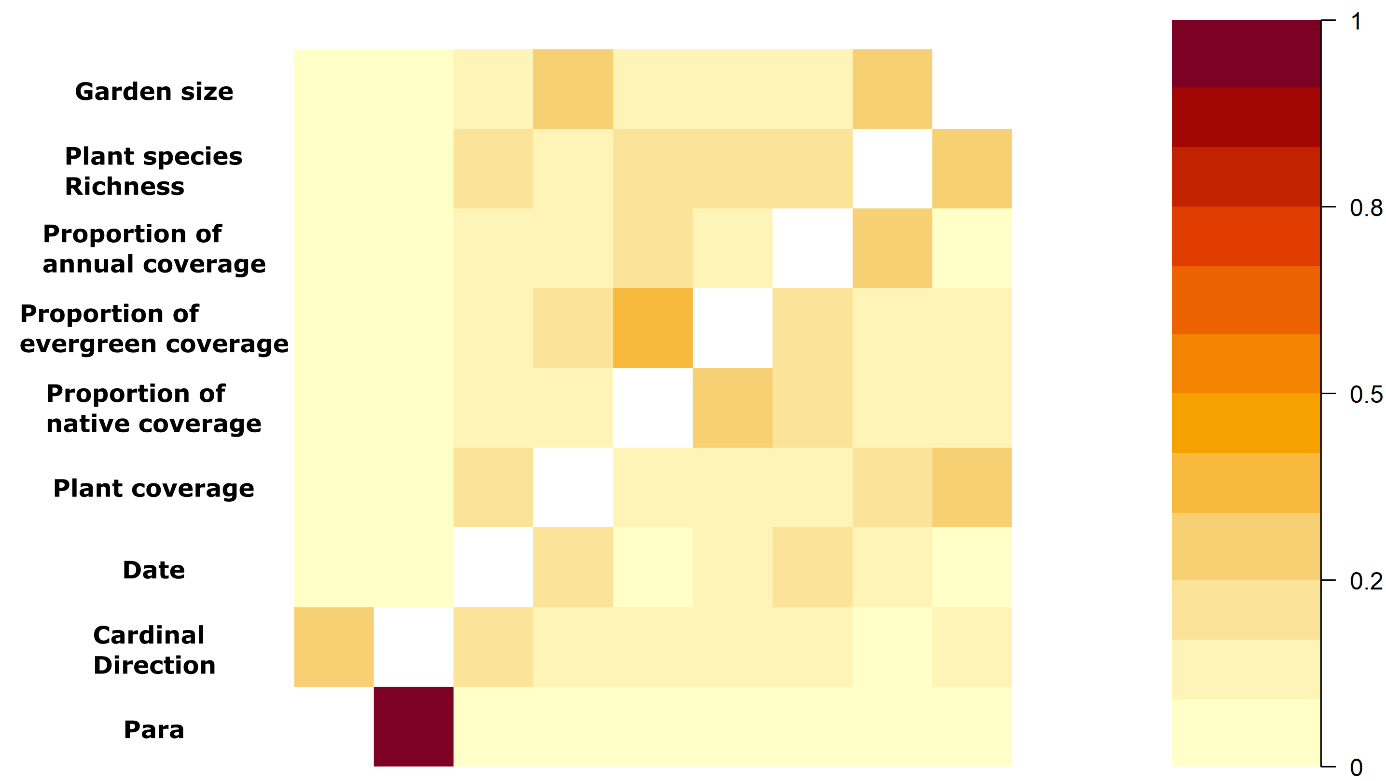

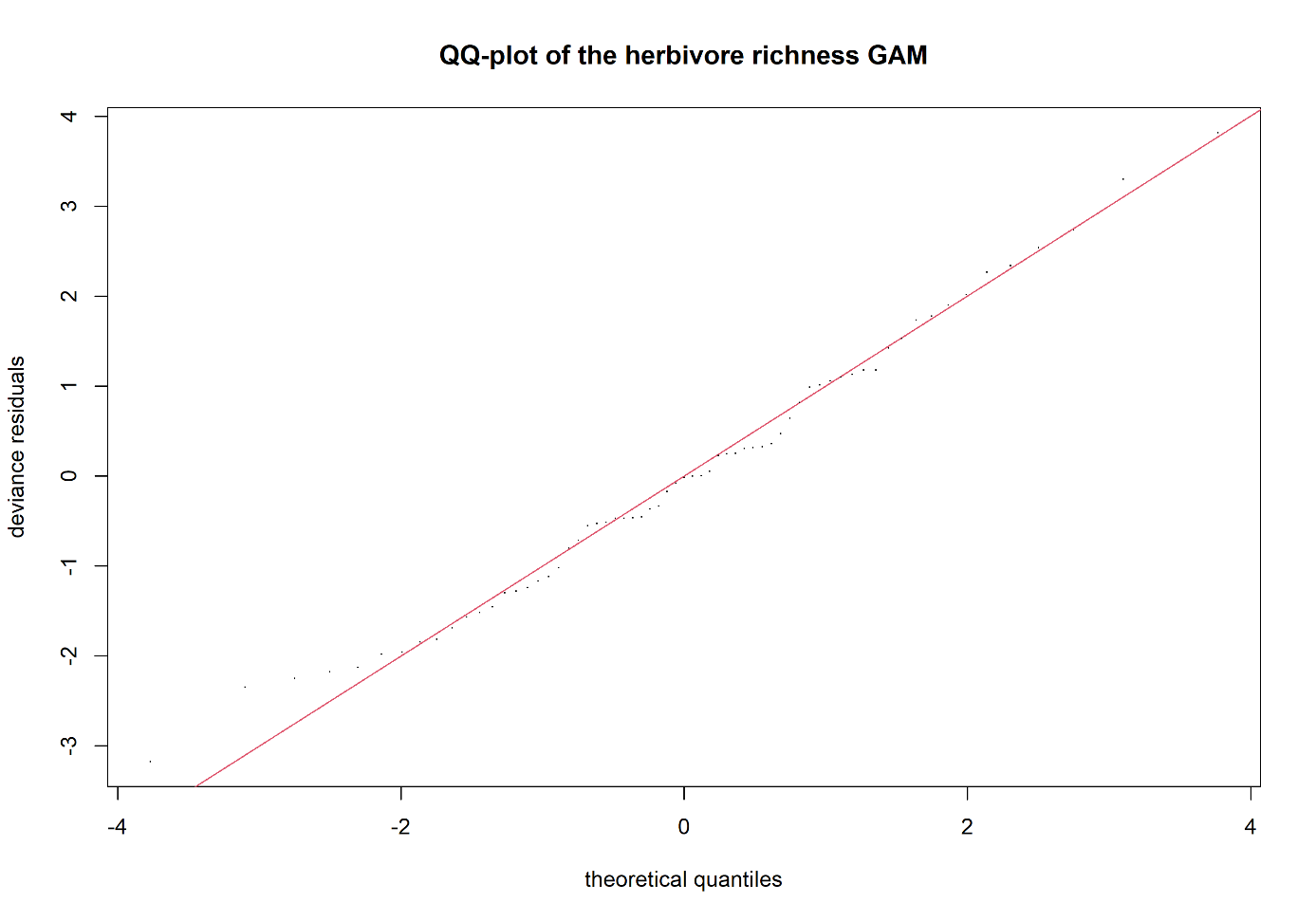

**Appendix 3. Shown are the model diagnostic plots for the herbivore species richness GAM covering ACF, concurvity and model fit.** None of the model diagnostics show any apparent problem with the model fit.

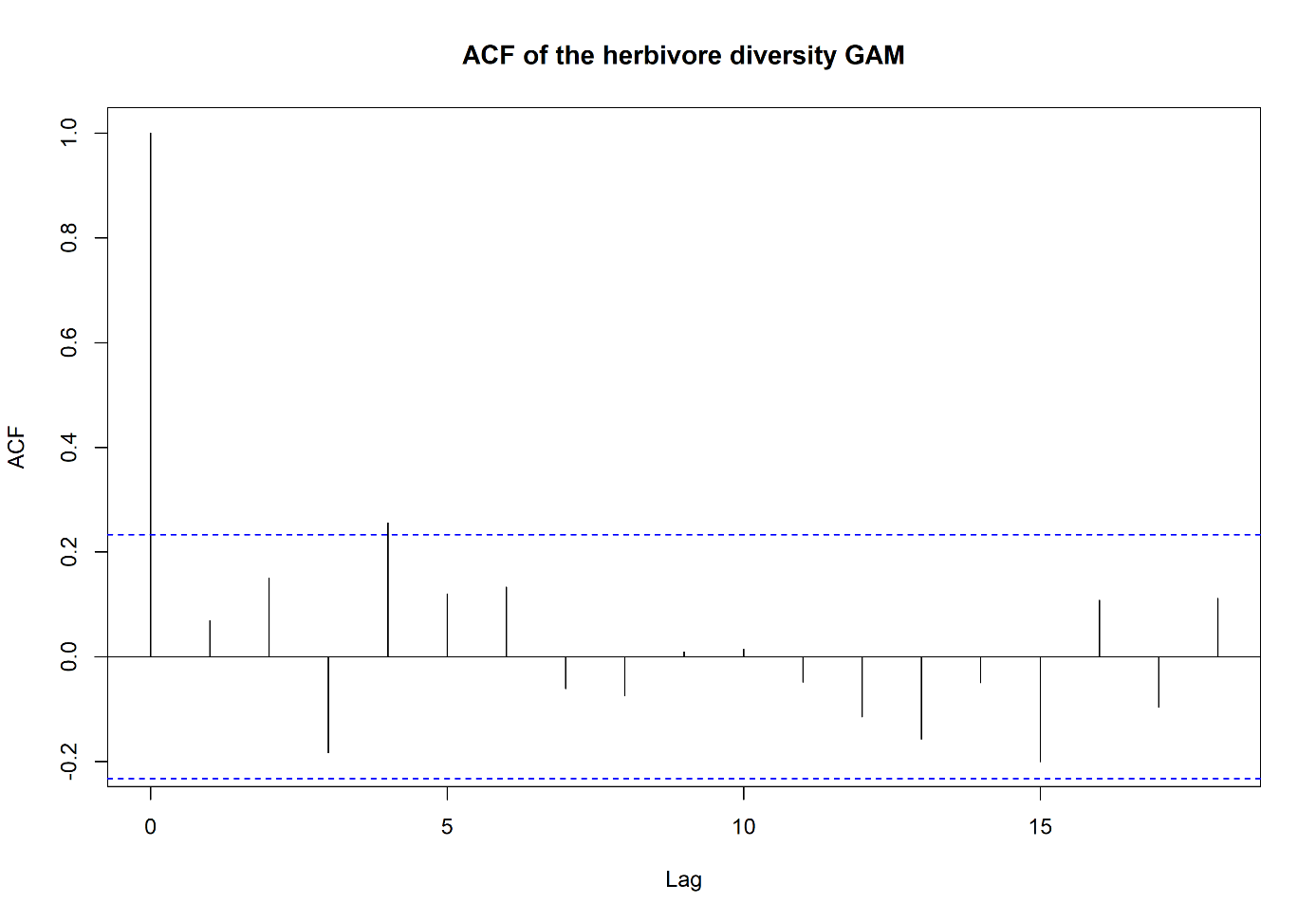

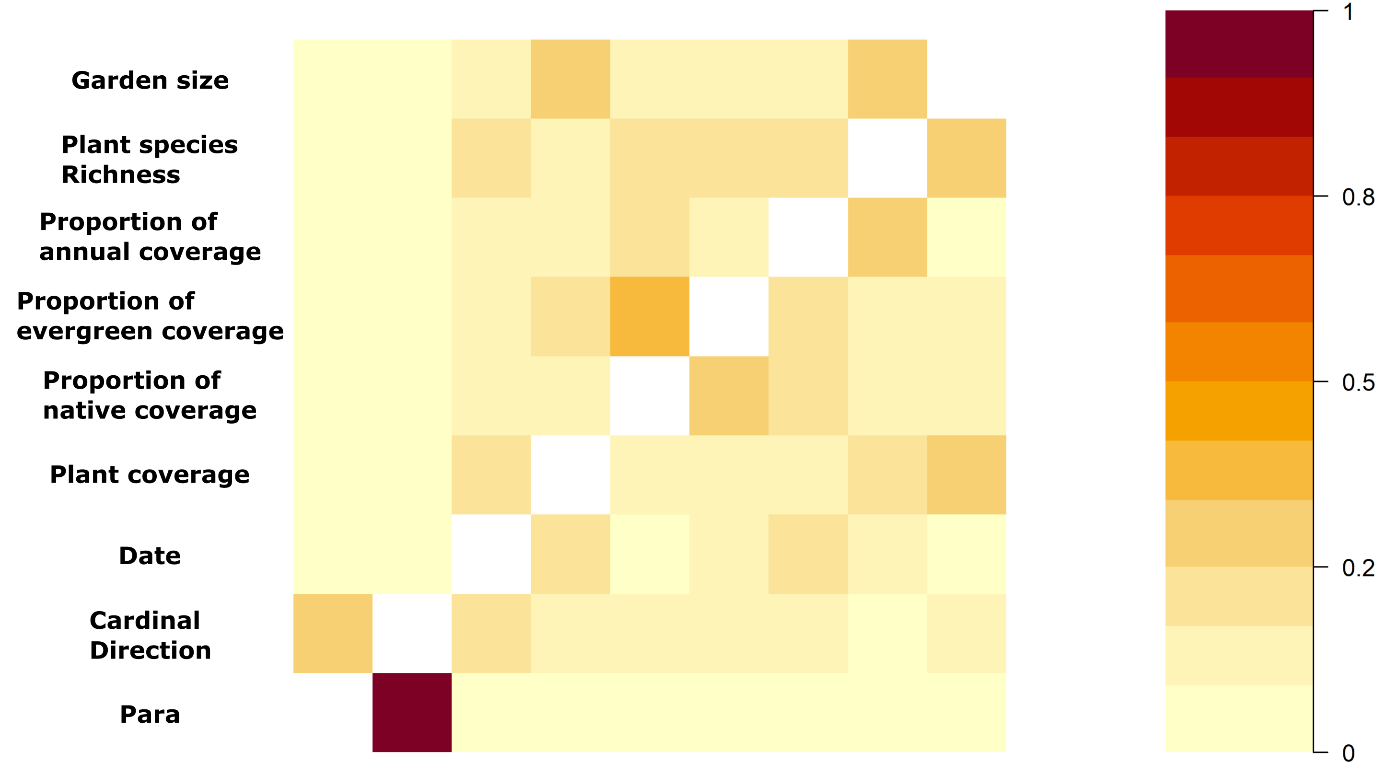

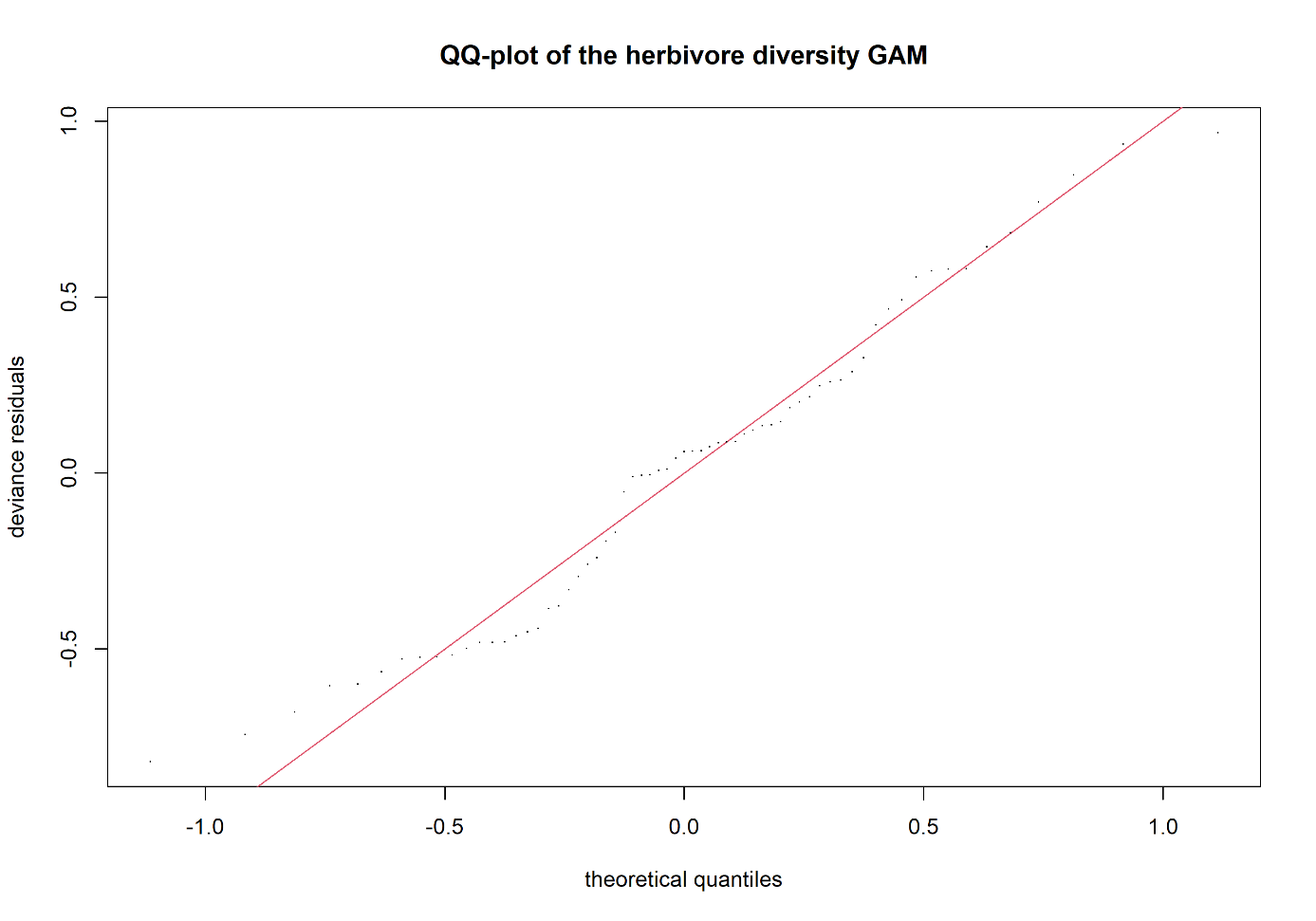

**Appendix 4. Shown are the model diagnostic plots for the herbivore diversity GAM covering ACF, concurvity and model fit.** None of the model diagnostics show any apparent problem with the model fit.
