## Supplementary material for "Vegetation density is the main driver of insect species richness and diversity in small private urban front gardens": Pollinator models

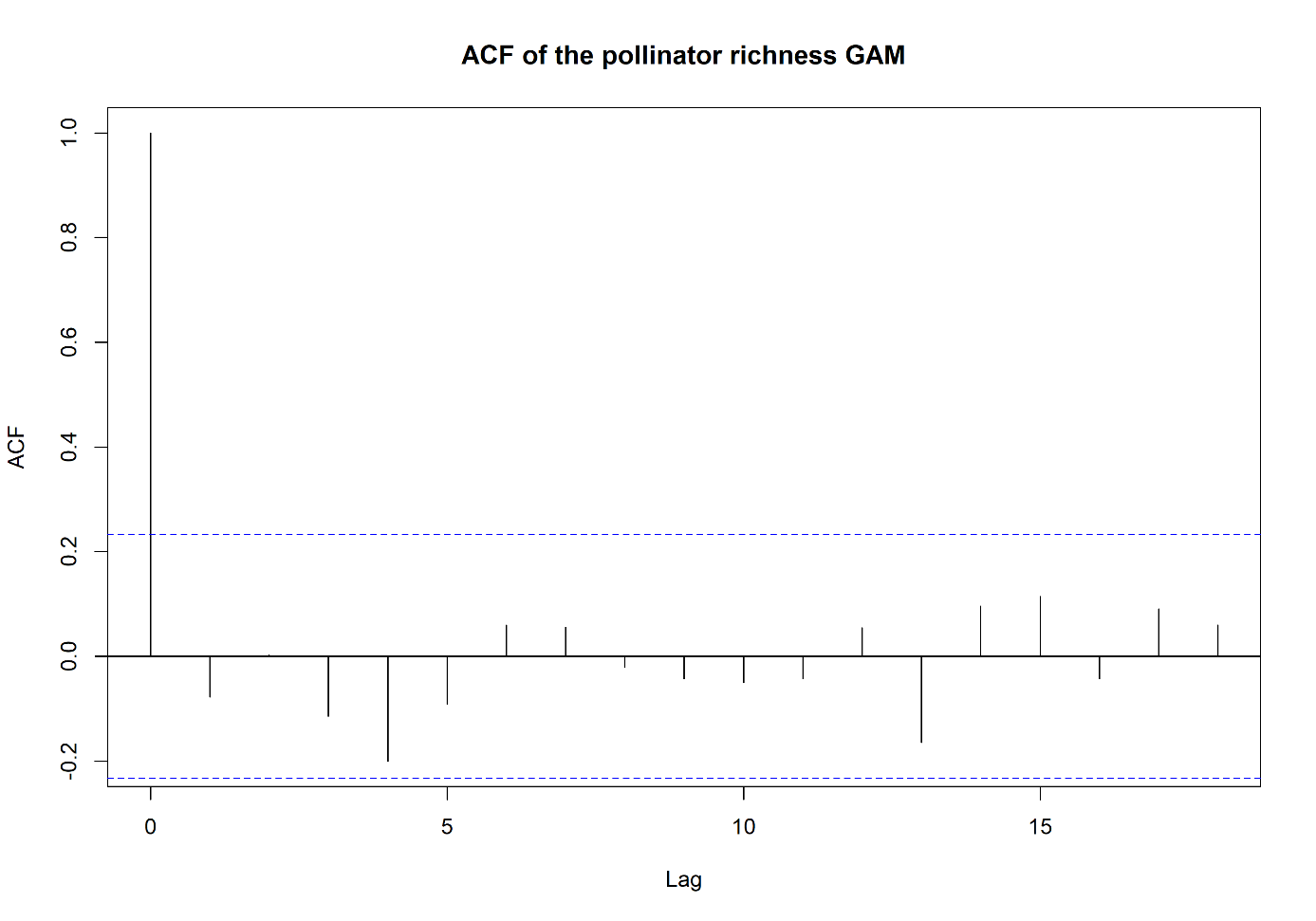

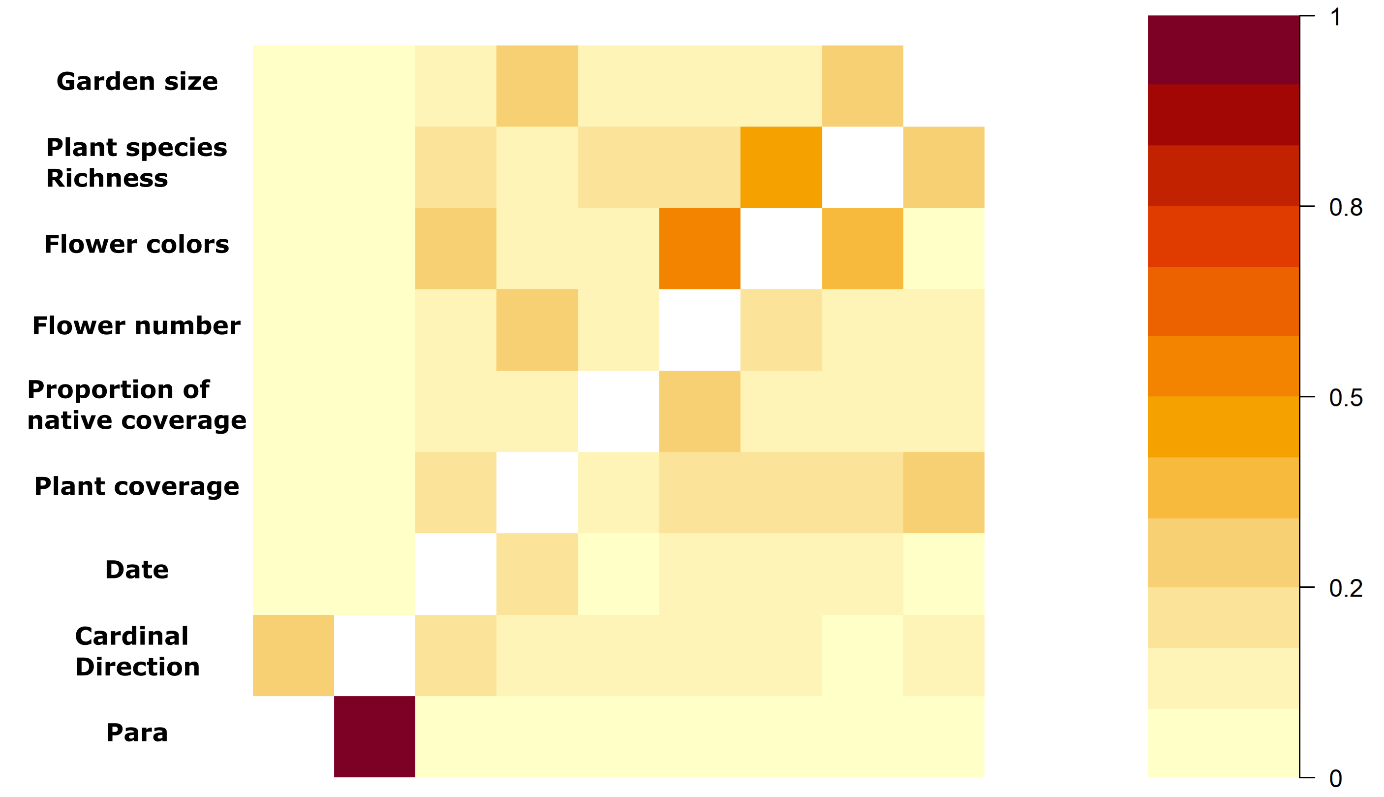

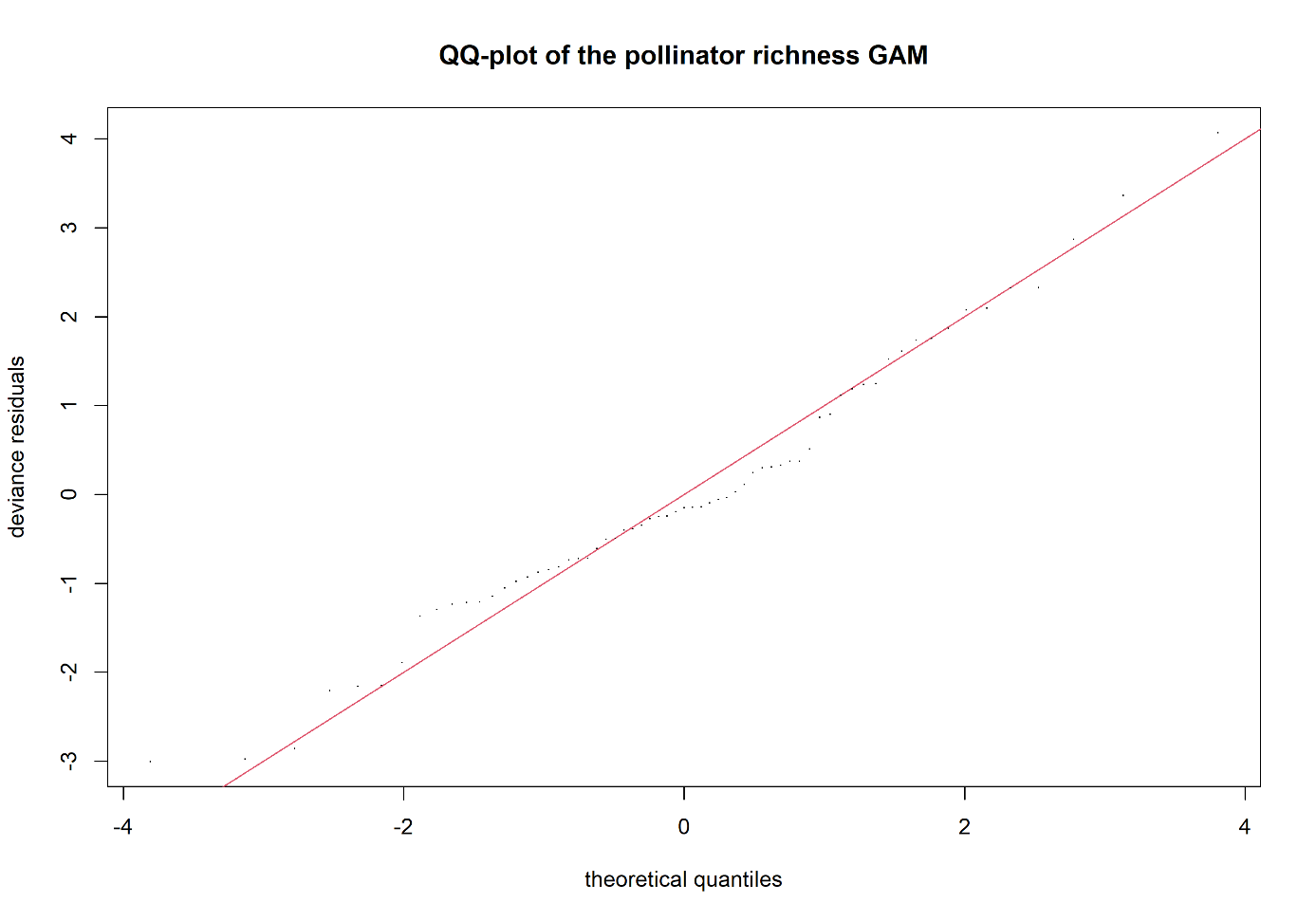

**Appendix 5. Shown are the model diagnostic plots for the pollinating species richness GAM covering ACF, concurvity and model fit.** None of the model diagnostics show any apparent problem with the model fit.

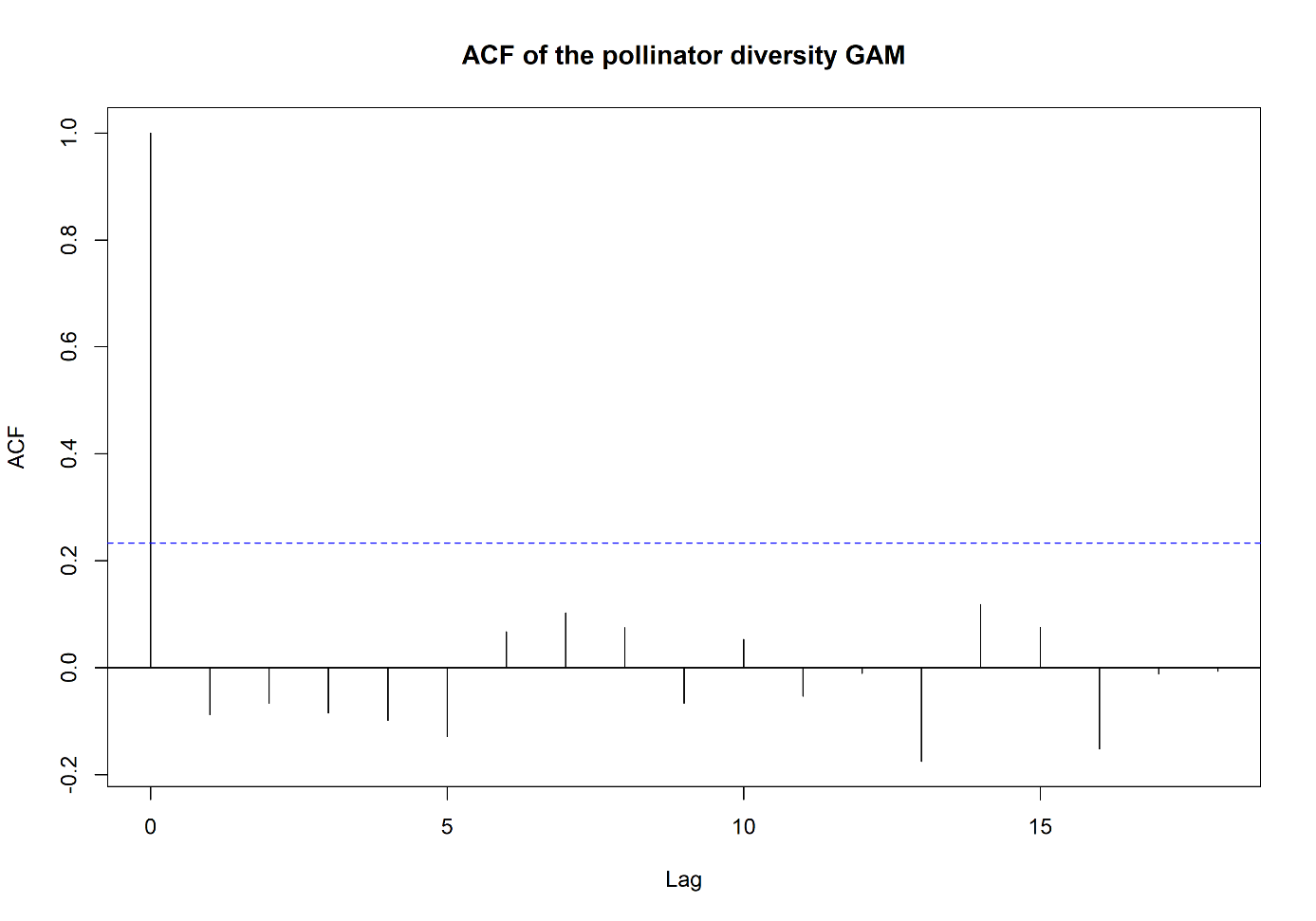

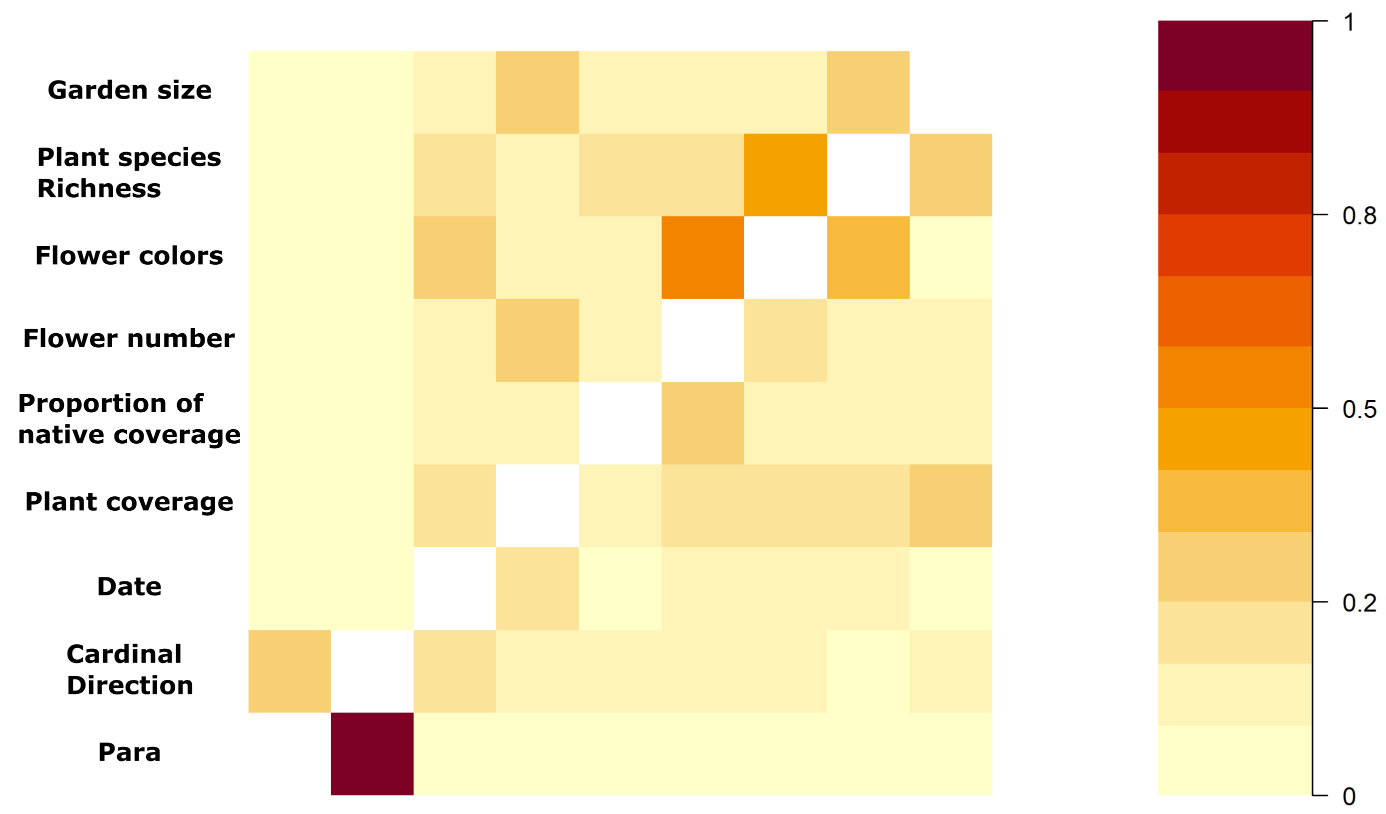

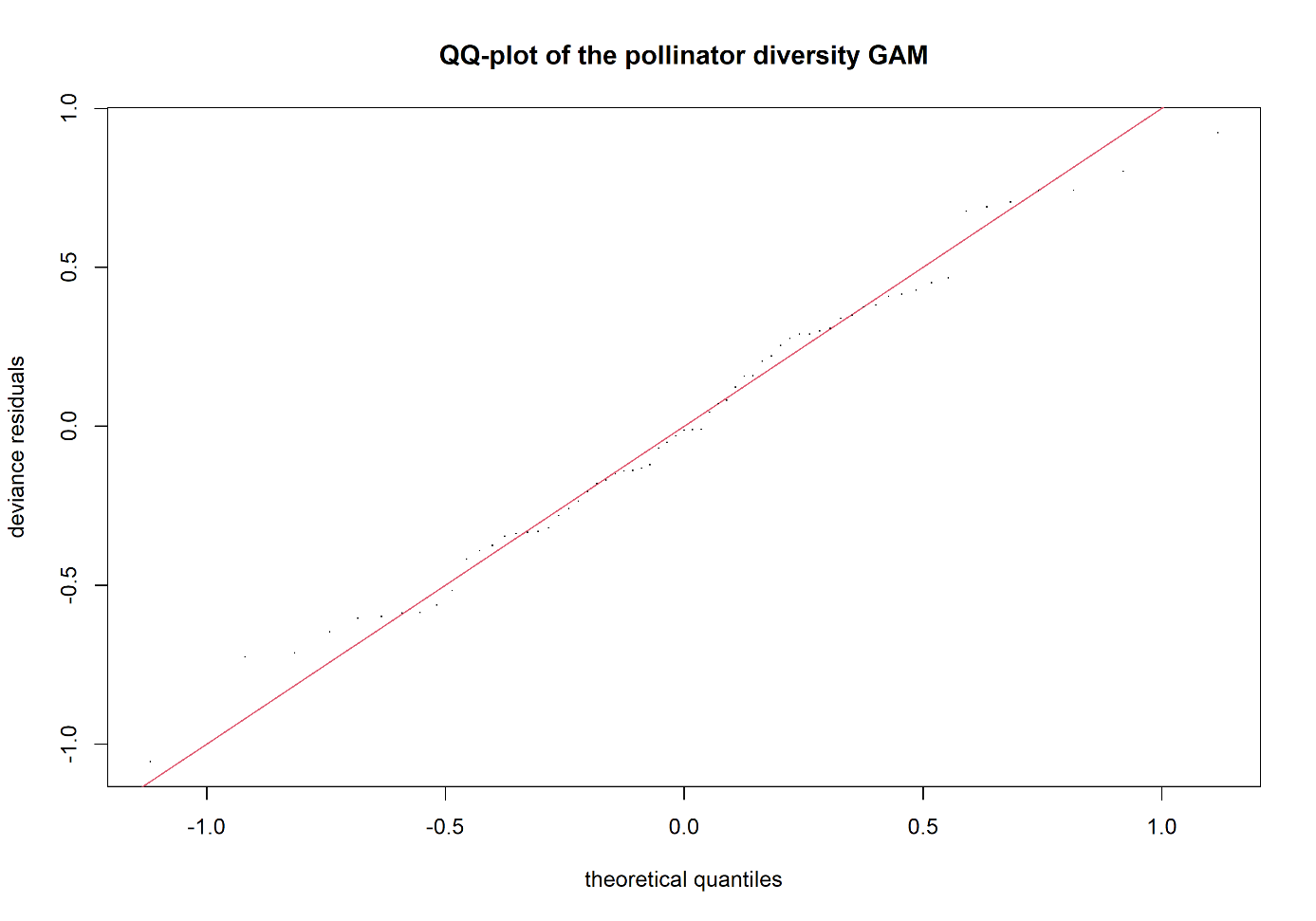

**Appendix 6. Shown are the model diagnostic plots for the pollinating diversity GAM covering ACF, concurvity and model fit.** The model diagnostics show a slight deviation in the ACF plot and some deviation from the red line in the QQ-plot.
