## Supplementary material for "Vegetation density is the main driver of insect species richness and diversity in small private urban front gardens": Insect models

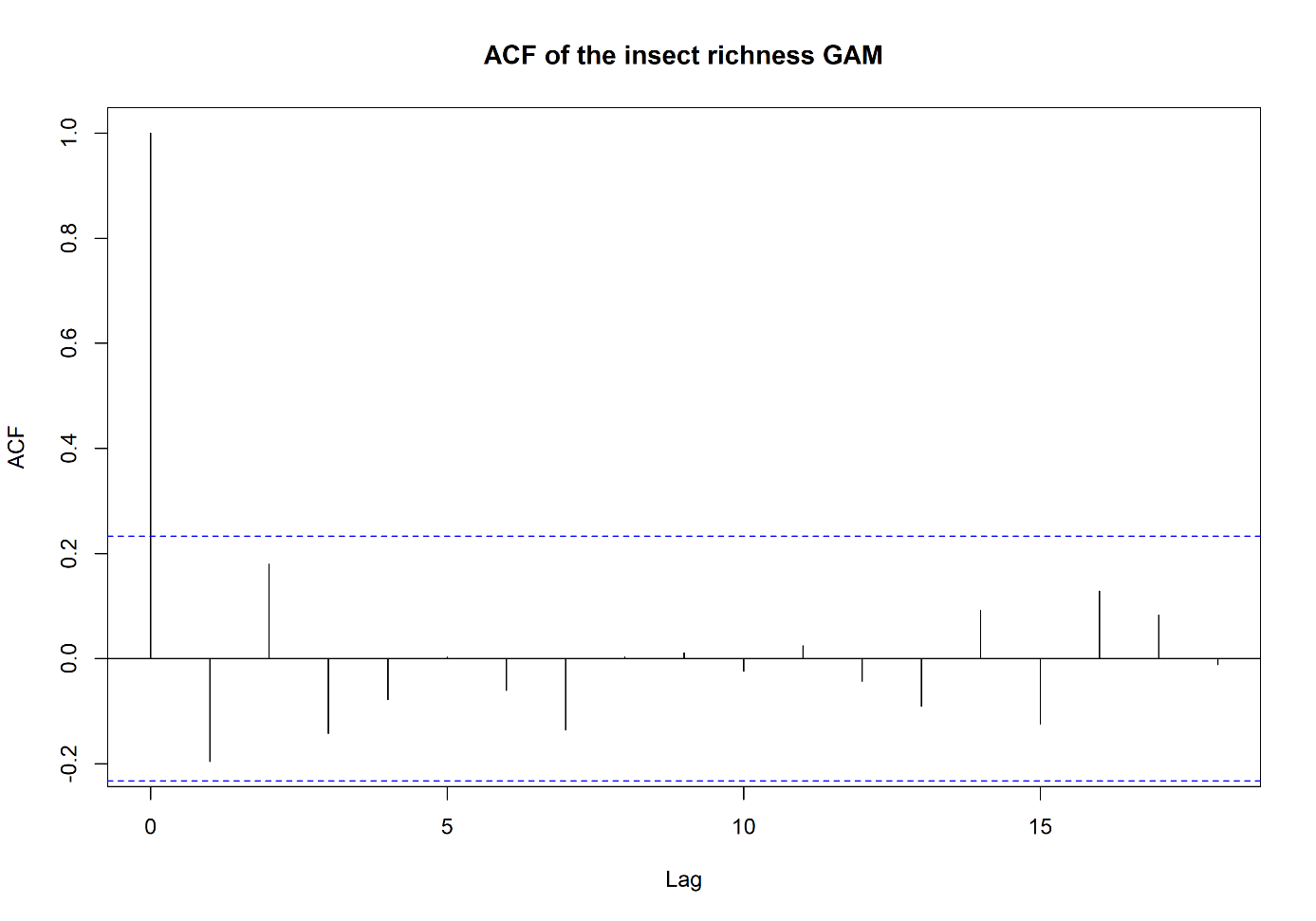

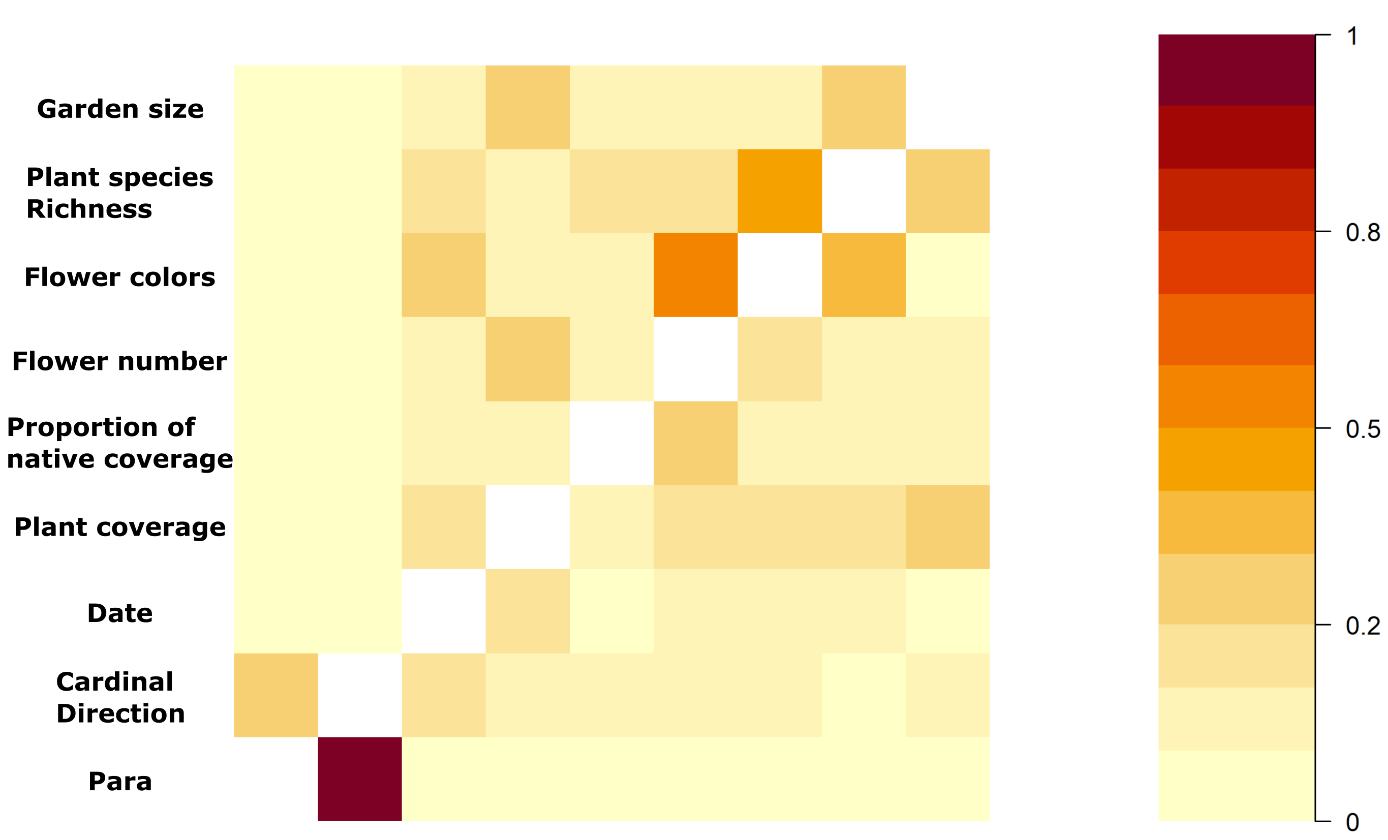

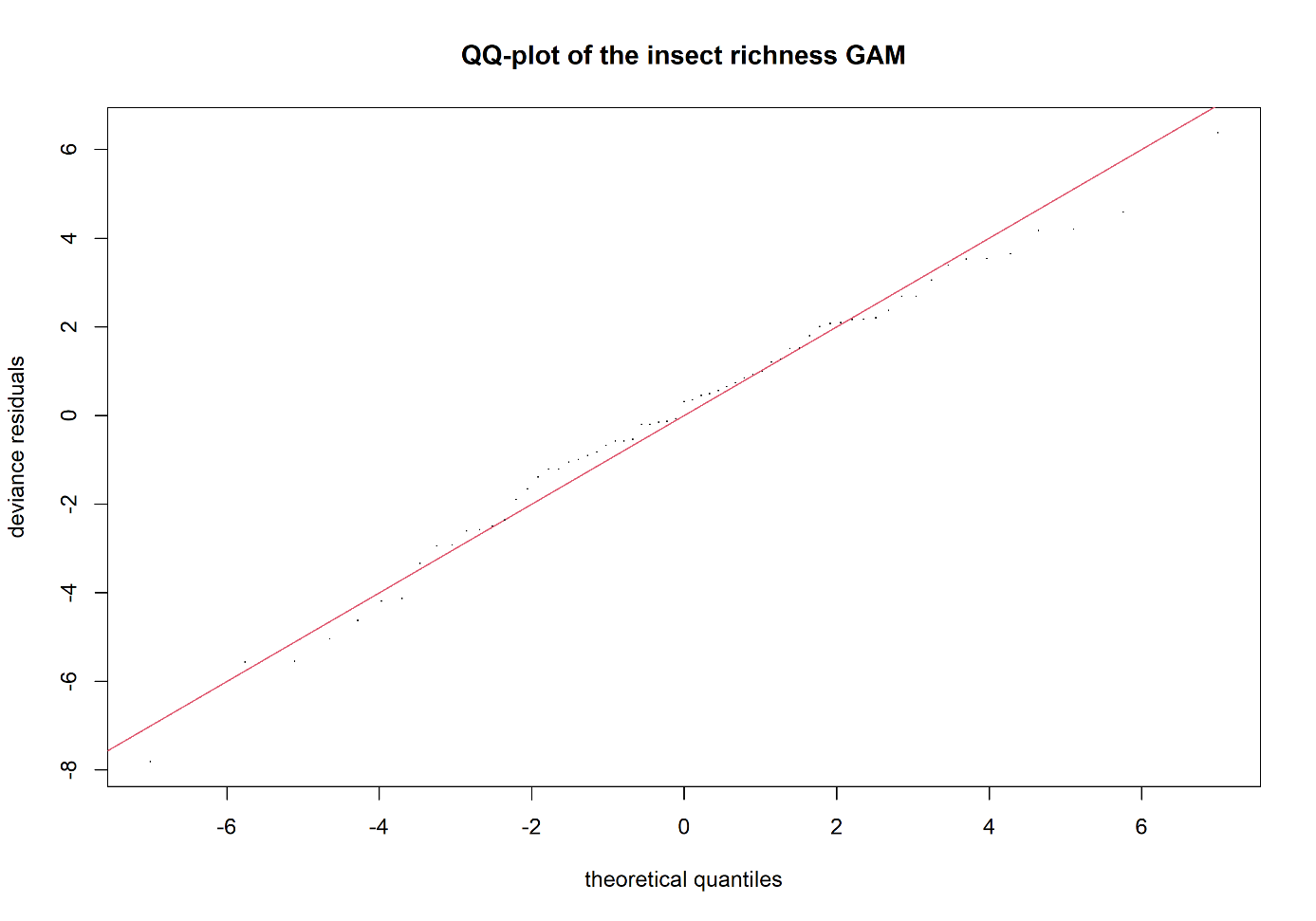

**Appendix 7. Shown are the model diagnostic plots for the insect species richness GAM covering ACF, concurvity and model fit.** None of the model diagnostics show any apparent problem with the model fit.

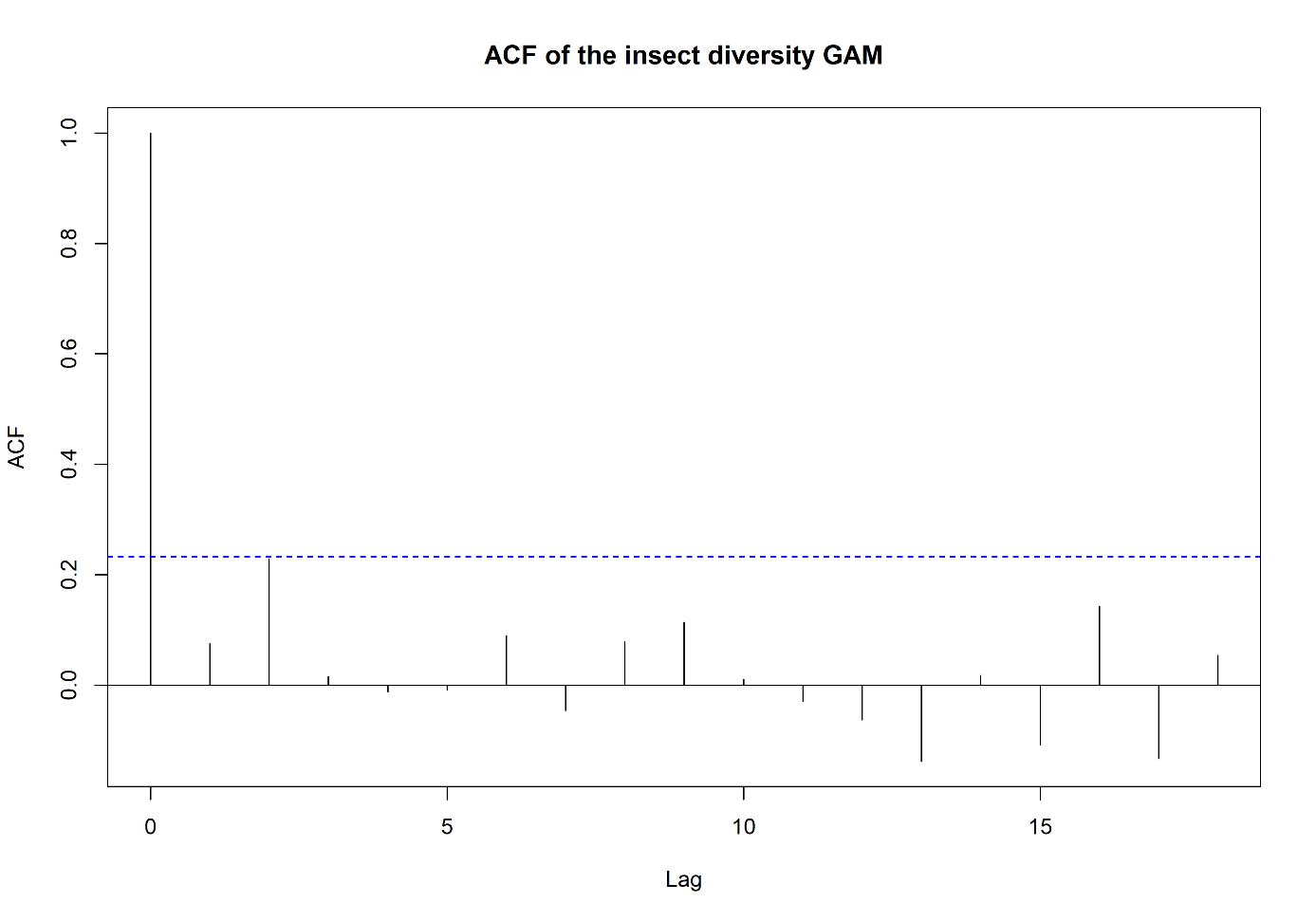

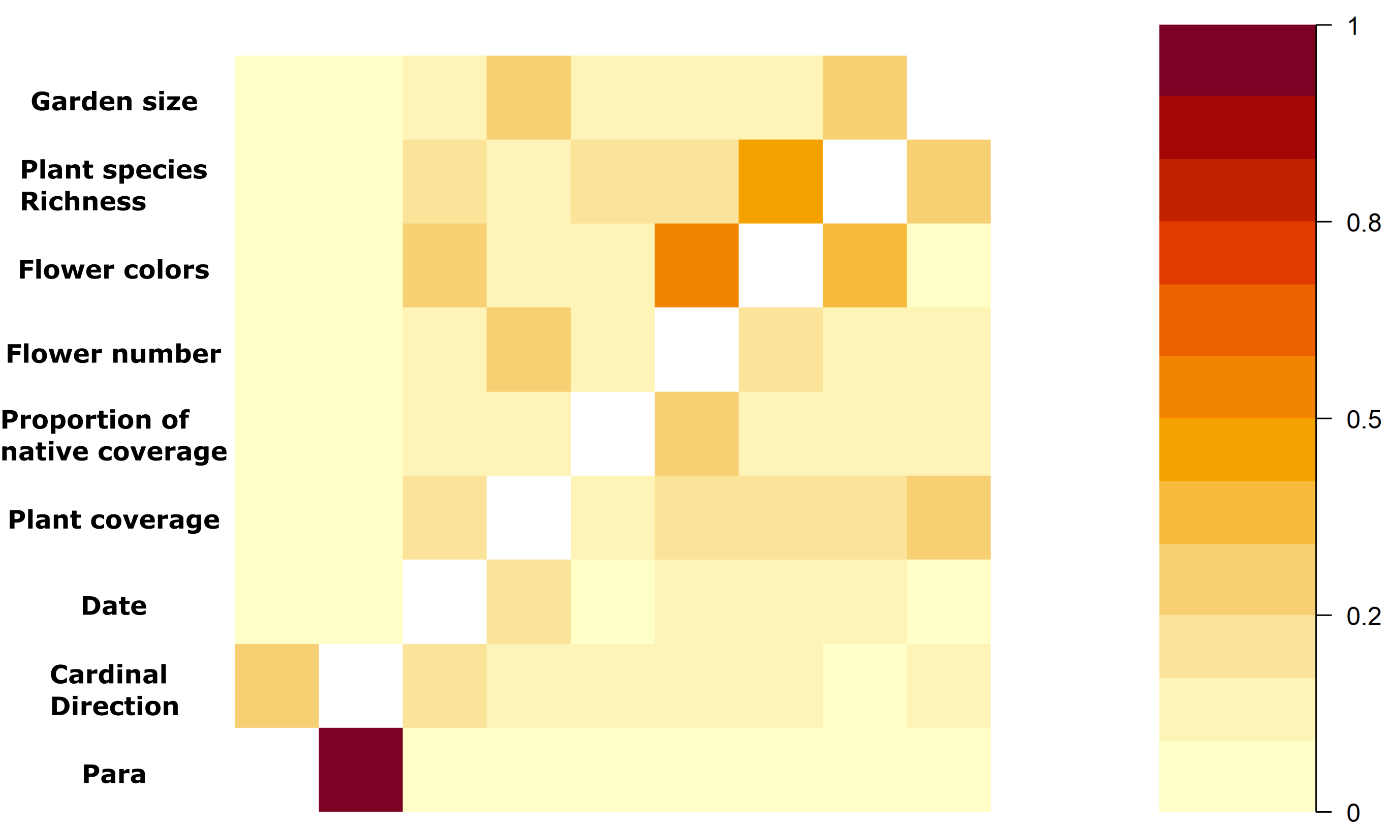

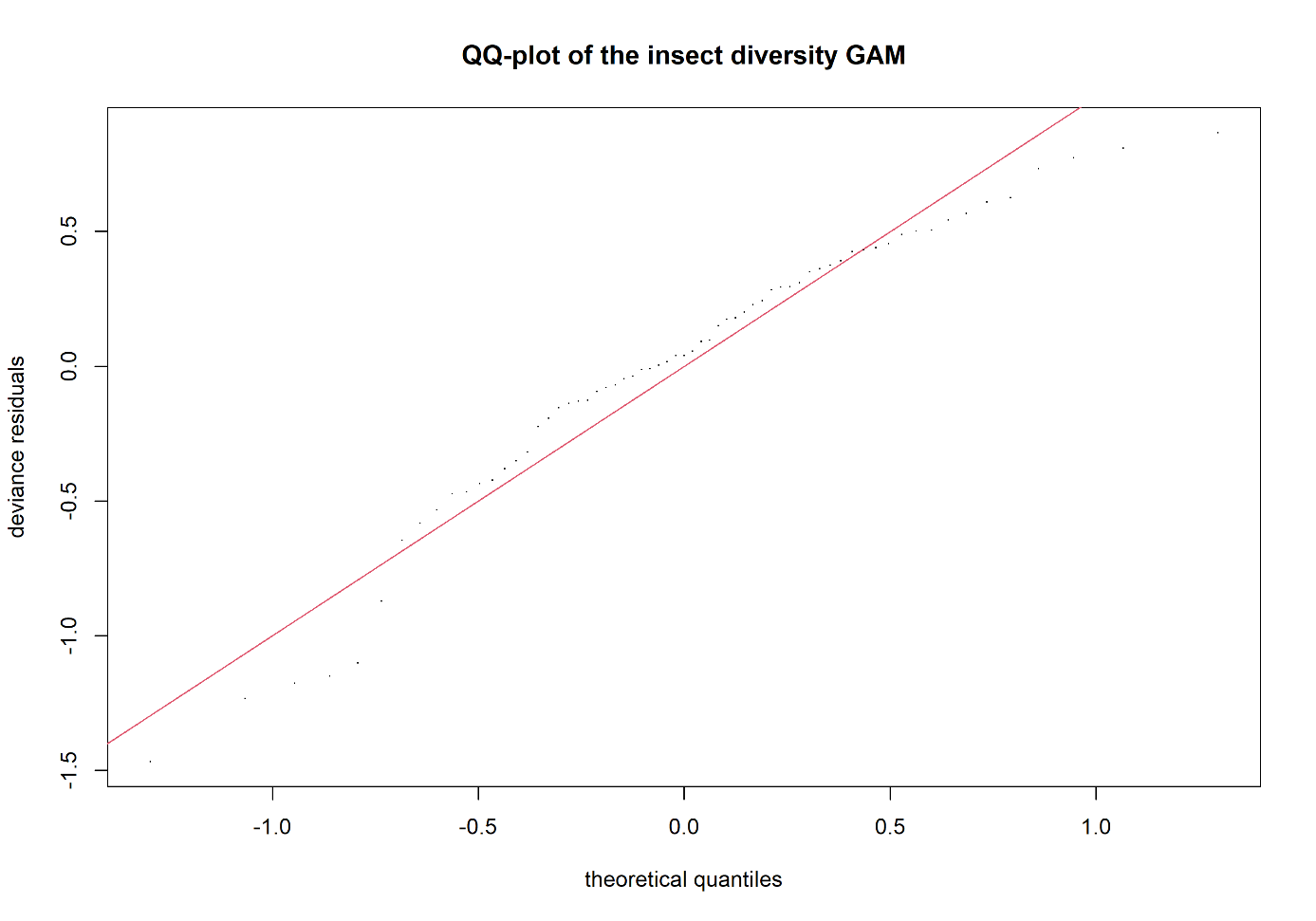

**Appendix 8. Shown are the model diagnostic plots for the insect diversity GAM covering ACF, concurvity and model fit.** None of the model diagnostics show any apparent problem with the model fit.
